# Activation of a cortical neurogenesis transcriptional program during NEUROD1-induced astrocyte-to-neuron conversion

**DOI:** 10.1101/2023.09.24.559157

**Authors:** Wen Li, Dan Su, Xining Li, Kang Lu, Qingpei Huang, Jiajun Zheng, Xiaopeng Luo, Gong Chen, Xiaoying Fan

## Abstract

NEUROD1-induced astrocyte-to-neuron (AtN) conversion has garnered significant attention as a potential therapeutic intervention to neurological disorders. To gain insight into the molecular regulations underlying this neuronal reprogramming process, we applied single-cell multiomics analyses on *in vitro* ND1-induced AtN conversion to systematically investigate how ND1 changed the fate of astrocytes at transcriptomic and epigenetic levels. Our findings reveal that the initial immature astrocytes go through an intermediate state where both astrocytic and neuronal genes are activated at early stage of AtN conversion. ND1 directly reshapes the chromatin accessibility landscape of astrocytes to that of neurons, promoting expression of endogenous *Neurod1 and other* neurogenic genes such as *Hes6, Insm1* etc. Interestingly, cell proliferation status is highly correlated with conversion rate, and inhibition of cell division significantly reduces the conversion ratio. Moreover, in comparison with another AtN reprogramming transcription factor, ASCL1, external ND1 can activate endogenous *Neurod1* and directly promote neuronal gene transcription; whereas external ASCL1 hardly activates endogenous *Ascl1,* leading to slower and inefficient conversion. Together, our studies demonstrate that *in vitro* AtN conversion mimics neurogenic transcriptional program in embryonic neurogenesis.

## Introduction

Adult mammalian neurogenesis is limited in terms of the number and region of occurrence (Kempermann et al., 2018; Lei et al., 2019). Therefore, strategies that promote endogenous neurogenesis hold great potential for therapeutic interventions to neurological disorders (Bocchi et al., 2022; Lei et al., 2019; Vasan et al., 2021). Unlike neurons, macroglial cells, including astrocytes and oligodendrocyte progenitors (OPCs), can reactivate and proliferate under conditions of injury and disease (Burda and Sofroniew, 2014). These reactive glia exhibit characteristics similar to neural stem cells (NSCs) and can give rise to a few immature neurons under specific circumstances (Magnusson et al., 2014; Robel et al., 2011; Tai et al., 2021; Zamboni et al., 2020). The latent neurogenic capacity of these cells can be enhanced through ectopic expression of pro-neuronal transcription factors (TFs), microRNAs, PTBP1 knockdown, or small molecules, both *in vitro* and *in vivo (Lei et al., 2019; Vasan et al., 2021)*. *In vivo* neuronal reprogramming from glial cells has shown therapeutic effects in animal models of various neurological diseases (Chen et al., 2020; Ge et al., 2020; Tai et al., 2021; Wu et al., 2020; Zheng et al., 2022).

NEUROD1 (ND1) is an important TF involved in embryonic and postnatal neuronal development (Gao et al., 2009; Hevner et al., 2006). It has been shown to rapidly and efficiently convert astrocytes into functional neurons, both *in vitro* and *in vivo (Guo et al., 2014)*. ND1-mediated AtN conversion has proven effective in treating focal ischemic stroke in rodent and non-human primate models (Chen et al., 2020; Ge et al., 2020; Tang et al., 2021). The addition of another TF, DLX2, could regenerate GABAergic projection neurons in the mouse striatum, extending the lifespan of Huntingtons transgenic mice (Wu et al., 2020). However, the ND1-mediated AtN conversion has recently been challenged, as researchers claiming that most “converted” neurons were pre-existing neurons (Wang et al., 2021). One significant reason behind this controversy is the lack of full understanding regarding the dynamic process and molecular regulations underlying the conversion of astrocytes into neurons by ND1.

Single-cell analyses are valuable tools for unraveling cellular heterogeneity and tracing intermediate cell states that may be masked in bulk analyses (Armand et al., 2021). They expand our knowledge of the dynamic processes involved in normal development and disease pathogenesis (Armand et al., 2021). However, the application of single-cell analyses to neuronal reprogramming, particularly *in vivo* reprogramming, is limited, partially due to technical difficulties in isolating target cells from adult brains (Giehrl-Schwab et al., 2022; Magnusson et al., 2020; Zhang et al., 2022). Nevertheless, the limited data from previous single-cell analyses have provided valuable insights into the potency of starting cells, intermediate states, and critical regulatory pathways in neuronal reprogramming (Magnusson et al., 2020; Zhang et al., 2022). Therefore, it is worthwhile to employ single-cell analyses to study ND1-mediated AtN conversion.

To achieve this objective, we have established an *in vitro* platform of ND1-induced neuronal reprogramming, which enables tracing the fate of starting cells and monitoring the entire conversion process without contamination of endogenous neurons. Then, single-cell multiomics sequencing, combined with time-lapse live imaging, is used to elucidate how astrocytes isolated from the young postnatal rat cortex are converted into neurons and unravel the underlying molecular regulations.

## Results

### ScRNA-seq captures diverse intermediate cell states in ND1-induced neuronal reprogramming

To investigate the cellular mechanisms underlying ND1-induced neuronal reprogramming from astrocytes, we established an *in vitro* trans-differentiation platform. Primary astrocytes were isolated from postnatal rat cerebral cortices and subcultured for 5 passages in the presence of 10% serum to reduce progenitor cells before transducing with a retrovirus carrying the *CAG::NeuroD1-IRES-EGFP* (ND1) construct (Figure 1A and S1). Retrovirus carrying the *CAG::EGFP* (GFP) construct acted as a control. The expression of ND1 was detected at 3 days post-infection (DPI, Figure S1E). Following ND1 expression, the astrocytes gradually changed their morphology and started expressing TUJ1 from 3 DPI (Figure S1F). The proportion of TUJ1^+^ cells increased over time (Figure 1B and S1F). By 14 DPI, approximately 80% of GFP^+^ cells had become TUJ1 positive. In contrast, the control group transduced with GFP alone showed negligible TUJ1 expression at all time points examined (Figure 1B, S1C-S1E). In addition to TUJ1, other markers indicative of immature and mature neurons, including DCX, NEUN, MAP2, and SV2, were also observed following ND1 transduction (Figure S1G-S1J). Subtype analysis revealed that over 90% of the converted neurons were excitatory neurons expressing vGLUT1, and among them, 60% expressed TBR1 and CTIP2, markers representing deeper-layer cortical neurons (Figure S1H and S1K). Furthermore, the converted neurons at 30 DPI acquired electrophysiological features similar to those of primary neurons isolated from rat E16.5 cortices (Figure S2A-S2H). They predominantly formed functional excitatory neuronal circuits, exhibiting frequencies and amplitudes comparable to those of E16.5 cortical primary neurons (Figure S2I-S2L). These results demonstrate a rapid and efficient *in vitro* ND1-induced neuronal reprogramming platform using rat astrocytes, similar to previous studies conducted with mouse or human primary astrocytes (Guo et al., 2014).

**Figure 1.**
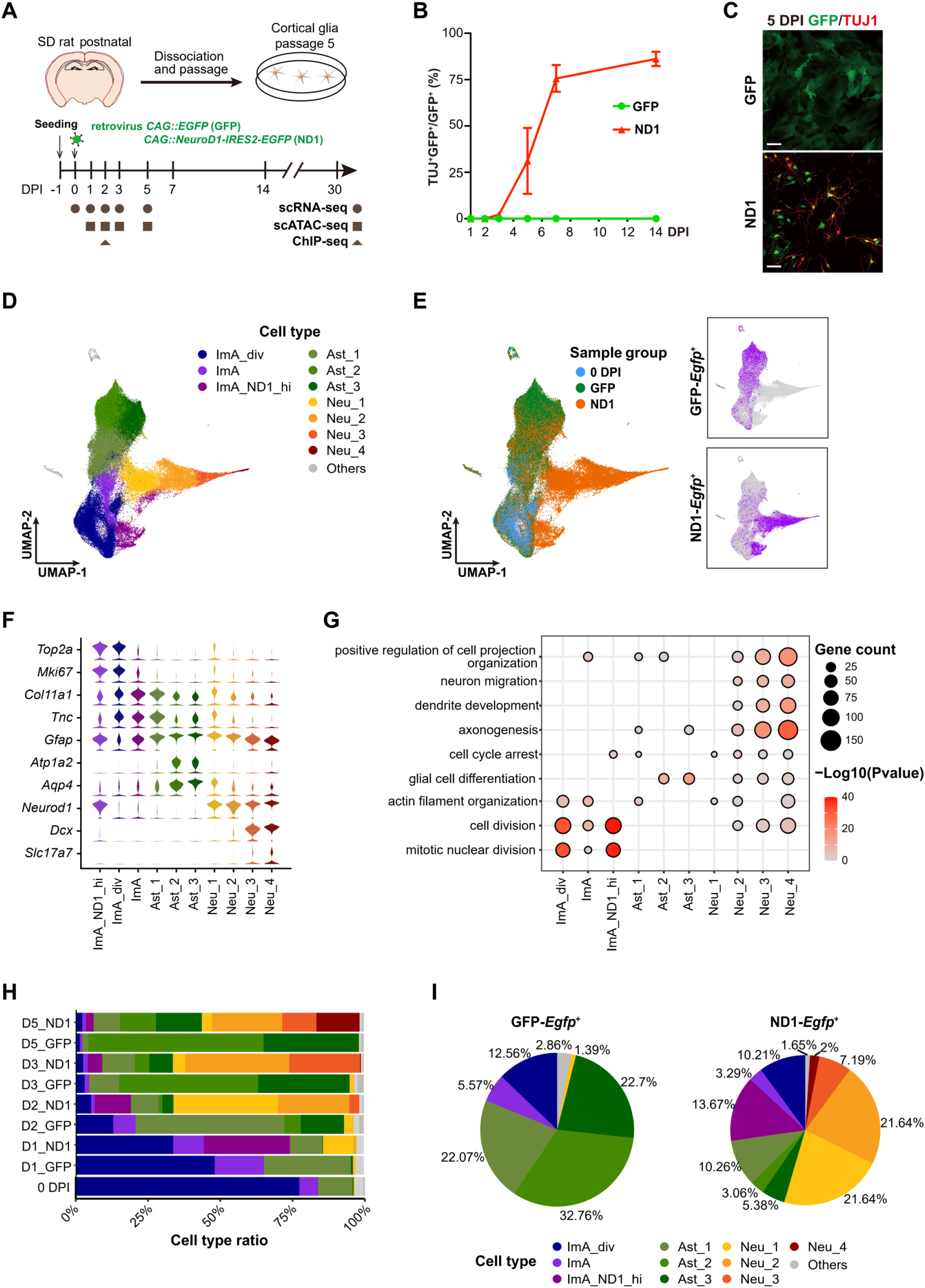
Cell type diversity in ND1-induced astrocytes to neuron conversion. (A) Schematic of setting up the *in vitro* AtN conversion system and sequencing strategies. (B) Quantitation revealing the increase in the ratio of converted neurons in ND1 group along time. (C) Representative imaging showing TUJ1^+^ (red) neurons at 5 DPI. Scale bar = 50 μm. (D) UMAP showing the cell types identified using scRNA-seq. (E) The group information of each cell. The *Egfp^+^* cells in each group are shown on the right. (F) The expression levels of classical markers in each cell type. (G) Representative GO terms of cell type DEGs. (H) The ratios of cell types in samples collected at indicated time points. (I) The ratios of cell types within the *Egfp^+^* cells in ND1 and GFP group.

Then, we utilized this platform to investigate the reprogramming mechanism through single-cell multiomics analyses (Figure 1A). Firstly, we generated a scRNA-seq dataset from cells from 0 to 5 DPI, when apparent fate conversion to neurons occurs (Figure 1C), to capture the dynamic cell types during the reprogramming process. After quality control, we retained 13,081 cells from 0 DPI (initial cells), 32,741 cells from the GFP control group, and 46,164 cells from the ND1 group for further clustering (Figure 1D, 1E). Based on a panel of known markers, we identified three main cell types: immature astrocytes (ImA), astrocytes, and neurons (Figure 1D, 1F, and Table S1). The immature astrocyte population could be further divided into three subtypes: ImA_div, ImA, and ImA_ND1_hi. They all exhibited high expression of *Gfap and Tnc* (Figure 1F). ImA_div cells showed high activity in the cell cycle, while ImA_ND1_hi shared many features with ImA_div but expressed high levels of *Neurod1* (Figure 1F). The astrocyte population consisted of three subclusters (Ast_1, Ast_2, Ast_3), with Ast_1 expressing high level of *Gfap, Tnc,* and *Col11a1,* while Ast_2 and Ast_3 expressing high level of *Atp1a2* and *Aqp4* (Figure 1F). The neuronal lineage comprised four subclusters (Neu_1, Neu_2, Neu_3, Neu_4) that exhibited high expression of neuronal markers and genes enriched in gene ontology (GO) terms related to axonogenesis, dendrite development, neuron migration, and positive regulation of cell projection (Figure 1F and 1G).

Notably, the starting cells (0 DPI initial cells, Figure S1) were predominantly identified as ImA immature astrocytes (approximately 80%, Figure 1H). The GFP control cells were primarily classified as immature astrocytes and astrocytes (Figure 1E and 1H), while the majority of cells in the ND1 group were clustered into the neuronal lineage (Figure 1E). Among the *Egfp*^+^ cells in the ND1 group, 32.2% were immature astrocytes and astrocytes, 13.7% were ImA_ND1_div, and 52.5% were neurons (Figure 1I). ImA_ND1_hi cells were exclusively found in the ND1 group, with the highest proportion observed at 1 DPI (Figure 1H). These findings indicate that scRNA-seq analysis captured the dramatic switch in cell identities from immature astrocytes to neurons induced by ND1.

### An intermediate state expressing both astrocytic and neuronal genes at early stage of ND1-induced neuronal reprogramming

We then conducted pseudotime analysis to uncover the lineage relationships. There were two distinct developmental branches originating from the immature astrocytes (Figure 2A and S3A). Based on the expression patterns of the astrocyte gene *Aqp4* and the neuron gene *Tubb3*, we defined these two branches as the astrocyte branch and the neuron branch, respectively (Figure 2B and S3A). Importantly, the sampled time points (0, 1, 2, 3, 5 DPI) aligned well with the pseudotemporal axis, indicating a high degree of consistency between the two (Figure 2A, 2C).

**Figure 2.**
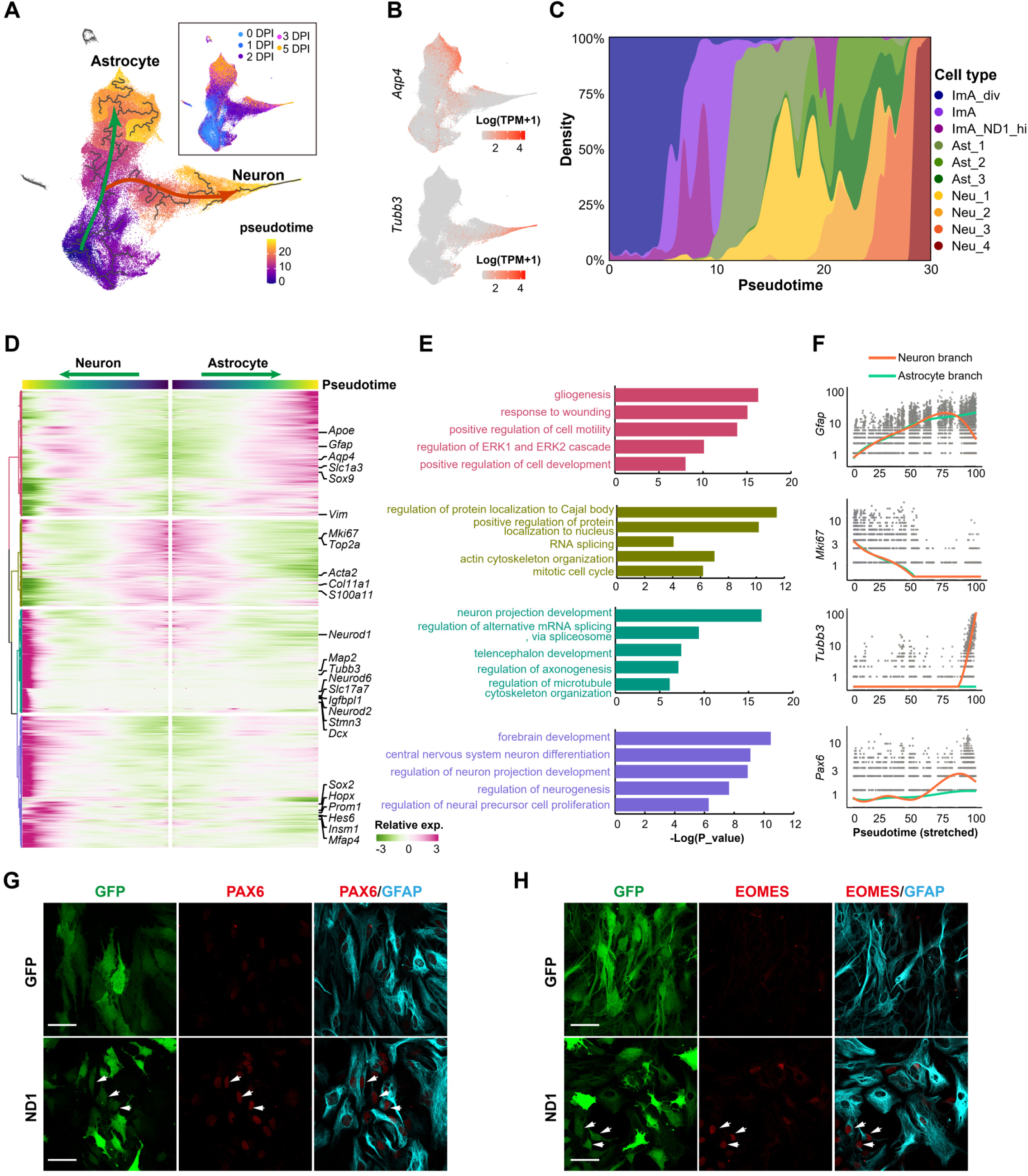
The trajectories of ImA to neuron reprogramming and to astrocyte development. (A) Pseudotime score of the cells on UMAP. Two branches are figured out. The real sampling time for the cells are shown in the right rectangle. (B) The expression of *Aqp4* and *Tubb3* are plotted to confirm the branches. (C) The density distribution of each cell type along the pseudotime. (D) Heatmap showing the expression patterns of the pseudotime genes along the two branches. Genes are clustered into four groups with different colors on the dendrogram. Representative genes in each group are also shown. (E) GO terms of genes corresponding to the groups in D. (F) Relative expression of representative genes corresponding to the four groups in D. (G-H) Representative images show the intermediate cells in the early stage of neuronal reprogramming (3 DPI), which co-express PAX6 (red) and GFAP (turquoise) (white arrows, G) or EOMES (red) and GFAP (turquoise) (white arrows, H). Scale bars = 50 μm.

To gain further insights into the molecular programs driving the conversion process, we extracted pseudotime genes along both trajectory branches and performed clustering analysis (Figure 2D, Table S1). In the astrocyte branch, immature astrocytes initially downregulated mitotic cell cycle genes (e.g., *Mik67* and *Top2a*, Figure 2D-2F and S3B), and subsequently upregulated genes involved in glial differentiation, including gliogenesis-related genes like *Gfap, Aap4,* and *Apoe*. On the other hand, immature astrocytes in the neuron branch also initially downregulated cell cycle genes and then underwent an intermediate state simultaneously upregulating genes associated with glial differentiation and early neuronal development, such as early neurogenesis genes like *Pax6* and *Sox2* (Figure 2D-2F and S3B). As the conversion proceeded, the intermediate cells dramatically decreased genes related to gliogenesis, and concomitantly increased genes associated with neuronal development events, including neuron projection development and regulation of neurogenesis (Figure 2D-2F and S3B). These findings suggest an important intermediate state that simultaneously upregulates both astrocytic and neuronal genes during the early stages of ND1-induced neuronal reprogramming of immature astrocytes.

Previous studies have suggested that ND1-induced AtN conversion bypasses the NSC stage (Guo et al., 2014), so we examined the expression levels of NSC markers *Sox2*, as well as markers of cortical neuronal progenitors, including Pax6, *Eomes* (*Tbr2*), and *Dcx (Hevner et al., 2006)*. Interestingly, *Sox2* and *Pax6* were transiently upregulated in the early stages of reprogramming and quickly downregulated (Figure 2F and S3B). Subsequently, *Eomes* and *Dcx* increased (Figure S3B). The immunostaining results supported the existence of PAX6^+^ and TBR2^+^ cells at 3 DPI in the GFP^+^ population of ND1 group (Figure 2G and 2H). These data suggest the presence of a transient stage of neuronal progenitor-like cells during the neuronal reprogramming of immature astrocytes.

To further elucidate the gene regulatory networks involved in the reprogramming process, we performed weighted correlation network analysis (WGCNA) based on all the cells and identified six gene modules closely associated with different cell types and states (Figure S3C, Table S1). The yellow module contains genes involved in cell division and exhibits a high score in the initial immature astrocytes (Figure S3D-S3F). The red module is shared by cells in the early stages of both astrocyte and neuronal lineages, enriched in genes participating in immune response (IL-7 response) and metabolism (ATP metabolic process and cell redox homeostasis). In contrast, the brown module is enriched in late-stage cells of both astrocyte and neuronal lineages, comprising genes involved in metabolic processes (response to nitrogen starvation and hormone metabolic process). The shift from the red module to the brown module along the pseudotemporal axis suggests a metabolic change during fate commitment and maturation (Figure S3D).

The astrocyte-associated network consists of two modules: the green module active along the astrocyte lineage and the turquoise module restricted to the late stage of astrocyte differentiation. Specifically, the green module is enriched in genes involved in regulating glial cell proliferation and the response to transforming growth factor beta (TGFβ) pathway, while the turquoise module contains genes that promote astrocyte differentiation and suppress neurogenesis processes (Figure S3D-S3F). The blue module comprises genes regulating neural progenitor proliferation and telencephalon development. The modules derived from WGCNA show high consistency with the previously identified gene clusters based on pseudotime analysis, further supporting the regulatory networks in ND1-induced neuronal reprogramming, which early activate the astrocyte and neuronal programs simultaneously, and then gradually shut down the astrocyte genes.

### Multiomics analysis identifies key downstream effectors in ND1-induced neuronal reprogramming

To understand how ND1 triggers neuronal transformation from immature astrocytes, we investigated the initial targets of ND1 binding. We performed NEUROD1 ChIP-seq for ND1-transfected cells at 2 DPI and identified 19,681 ND1 binding peaks (Figure S4A, Table S2). Most of the genes associated with these peaks were upregulated in the neuronal lineage (Figure S4B, Table S2), indicating that ND1 positively regulates these genes. Meanwhile, there were also some genes that were downregulated in the neuronal lineage upon ND1 binding, reflecting negative regulation by ND1. Gene ontology (GO) analysis revealed that the upregulated ND1 targets, such as *Hes6*, *Insm1*, *Id2*, *Dcx*, and *Tubb3*, are associated with neuronal development and differentiation (Figure 3A and S4C). On the other hand, the downregulated ND1 targets, including *Cebpd*, *Fos*, *Anxa1*, are mainly involved in the mitotic cell cycle and functions related to reactive astrocytes (such as response to wounding and extracellular matrix organization, Figure 3A, S4C). As expected, ND1 self-regulates its expression by directly binding to its own promoter (Pataskar et al., 2016). These findings suggest that ND1 can directly regulate gene expression in both positive and negative ways, driving immature astrocytes towards the neuronal lineage.

**Figure 3.**
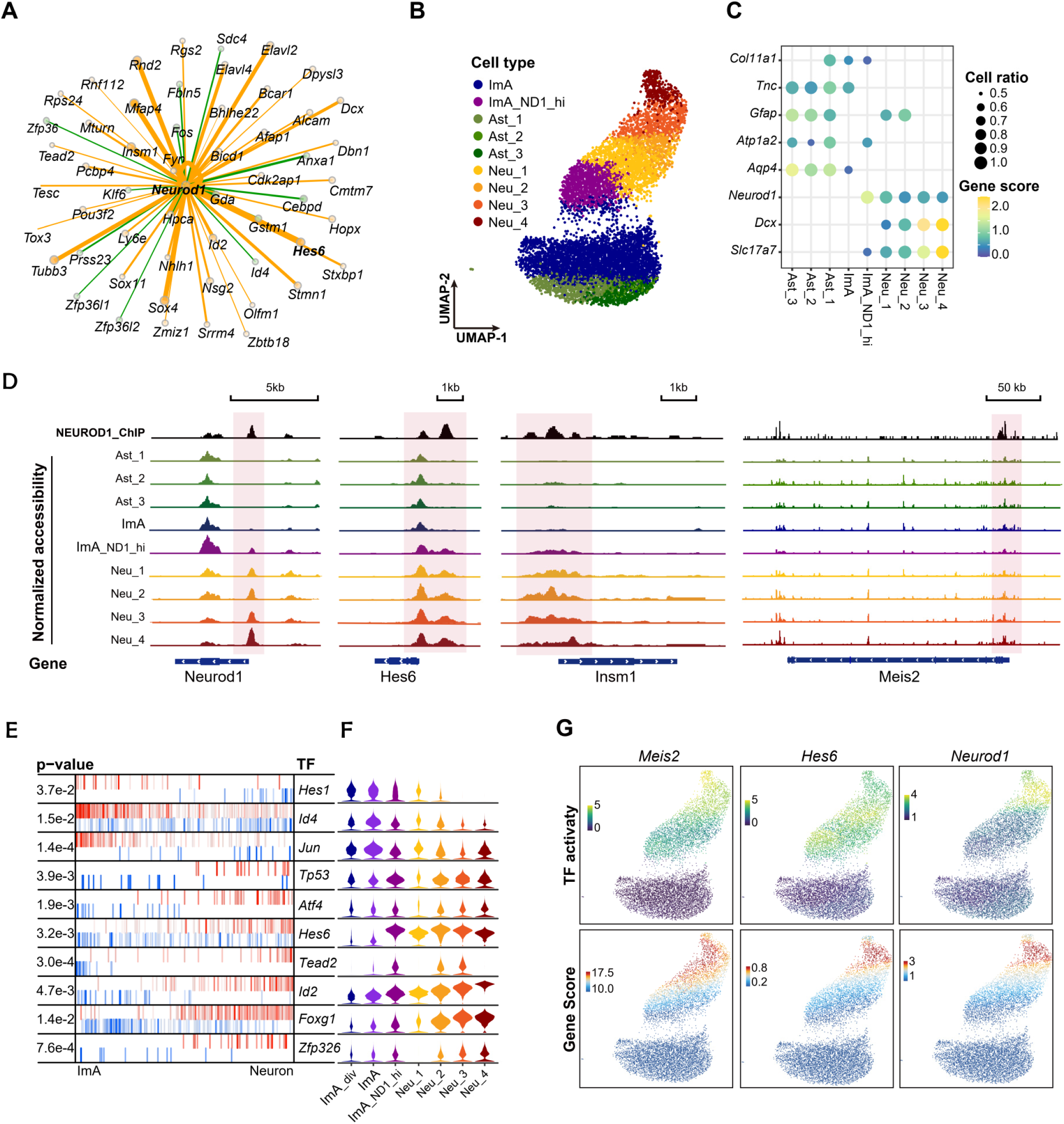
Important ND1 target TFs in neural reprogramming. (A) The top target genes of ND1 according to ChIP-seq results. The yellow line indicates ND1 positively regulated targets and the green line indicates ND1 negatively regulated targets. (B) UMAP showing the cell types identified using scATAC-seq. (C) Gene score of the classical markers in each cell type. (D) The genome browser view showing the NEUROD1 ChIP signal and chromatin accessibility signal. (E) Top 10 transcription factors regulating the reprogramming from ImA to neuron. The blue sticks represent repressed targets while the red sticks represent the activated targets. (F) The expression of the TFs indicated in E. (G) Gene score and TF activity predicted using scATAC-seq data.

To investigate the chromatin remodeling promoted by ND1 during neuronal programming, we performed scATAC-seq on cells from the ND1 group (Figure 1A). All cell types and subtypes identified in the scRNA-seq data were classified, except for ImA_div cells, as the cell cycle characteristics are masked in scATAC-seq data (Hsiung et al., 2015) (Figure 3B). Interestingly, the chromatin accessibility landscapes of immature astrocyte and astrocyte subtypes were highly similar to each other but distinct from the neuronal subtypes (Figure 3B). The gene scores of cell type marker genes validated the cell type annotation of the scATAC-seq data (Figure 3C and S4D). Importantly, Neu_1 and Neu_2 clusters showed high gene scores of both astrocytic gene *Gfap* and neuronal gene *Dcx*, supporting the transitional stage of converting cells in-between astrocytes and neurons during conversion process (Figure 3C). Genome browser visualization confirmed the specific opening status of cell type marker regions (Figure S4E). For example, *Aqp4* peaks were highly detectable in astrocyte lineages, while *Dcx* peaks were only present in neuronal lineages. *Gfap* peaks were observed in astrocytes and early reprogrammed neuronal subtypes but were not detected in the late neuronal subtype (Neu_4). When focusing on the chromatin accessibility of ND1-occupied loci, we found that these loci were more open in the neuronal subtypes (Figure S4F), indicating that ND1 enhances the accessibility of neuronal genes. Specifically, ND1-occupied and upregulated genes, such as *Neurod1*, *Insm1*, *Mesi2*, and *Hes6*, exhibited increased chromatin accessibility in neuronal cells (Figure 3D). Conversely, the downregulated ND1 targets, such as *Cebpd*, *Fos*, and *Anxa1*, gradually exhibited closed chromatin accessibility (Figure S4G). Collectively, these findings suggest that ND1 can directly bind to specific loci and rapidly induce chromatin remodeling, favoring the conversion from astrocytes to neurons.

To identify downstream regulators, we performed motif analysis of the scATAC-seq data and predicted regulatory TFs (Table S2). TFs associated with forebrain development and neuronal differentiation exhibited gradually increasing activity in the neuronal subtypes, while those associated with cell proliferation and astrocyte metabolic processes displayed decreasing activity (Figure S4H). To further identify key regulators, we extracted the most significantly regulated TFs along the AtN trajectory based on the scRNA-seq data. Examples include *Hes6, Id2, Tead2,* and *Id4*, which were directly bound by ND1 and showed upregulation of both themselves and their putative target genes along the neuronal lineage (Figure 3A, 3E and 3F). Interestingly, we identified opposite regulations in certain TF families during AtN conversion, such as a downregulation of Hes1 versus an upregulation of Hes6, and a downregulation of Id4 versus an upregulation of Id2 (Fig. 3F). These results suggest that different TFs within the same family may play opposite functions in cell fate conversion. The scATAC-seq results also revealed increased gene scores and motif activities for core TFs such as *Neurod1, Hes6*, and *Mesi2* along the cell fate transition (Figure 3D and 3G), further indicating their importance in ND1-induced neuronal reprogramming (Masserdotti et al., 2015; Matsuda et al., 2019).

### ND1-induced neuronal reprogramming partly resembles the cortical deeper-layer neurogenesis

In the developing cortex, ND1 has been identified as a critical transcription factor involved in the specification of deeper-layer neurons (Hevner et al., 2006). In this study, the majority of ND1-reprogrammed neurons were identified as cortical excitatory neurons (Figure S1H and S1K). We wondered whether *in vitro* reprogramming and *in vivo* cortical development share similar cellular and molecular processes.

To address this question, we first profiled cells from E16.5 and P2 rat cerebral cortices using scRNA-seq (Figure 4A and S5A-S5D, Table S1). We found that deeper-layer neurons were predominantly captured from E16.5, while upper-layer neurons were mainly detected at P2. The cell composition was further confirmed by immunostaining (Figure S5C and S5D). We then compared the *in vitro* reprogrammed cells with the cells from *in vivo* cortical development. As expected, the immature astrocytes and Ast_1,2,3 cell populations were assigned to astrocytes *in vivo* (Figure 4B and S5E). The Neu_1 cells were primarily aligned with astrocytes *in vivo*, and the subsequent neuronal subtypes exhibited a gradual transition from astrocytes to Ex_DL_3 through the radial glia (RG) and intermediate progenitor cell (IPC) states (Figure 4B and S5E). Immunostaining of the cells at 5 DPI indeed showed the presence of CTIP2^+^ neurons (Figure 4C). These findings suggest that ND1 might induce a trans-differentiation program that mimics the neurogenic paradigms of *in vivo* neurogenesis, progressing from RG to IPCs and then to deeper-layer neurons (Figure 2G and 2H).

**Figure 4.**
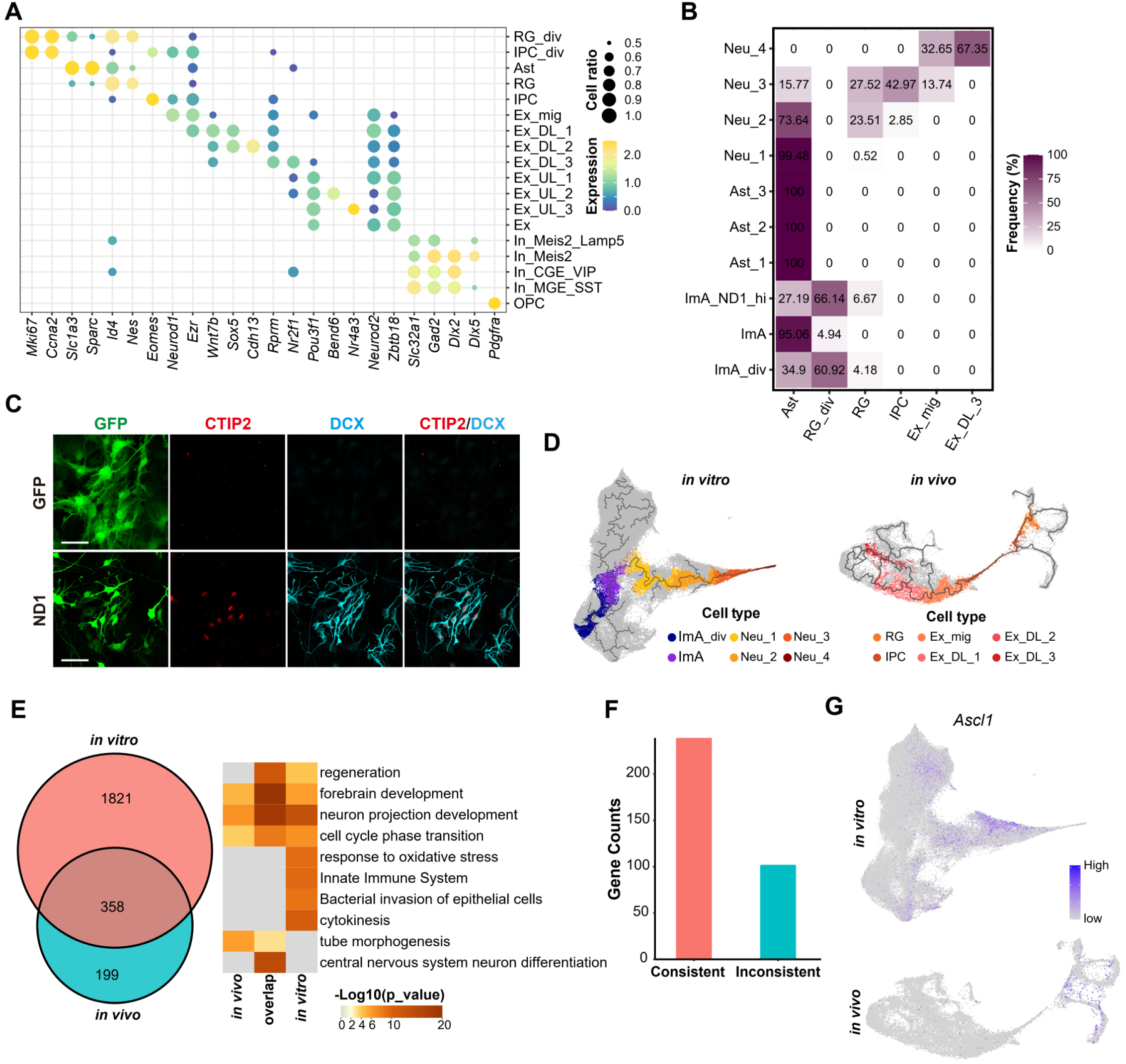
Comparison of ND1-induced neuronal reprogramming to the in vivo neurogenesis. (A) Expression levels of the markers in each cell type identified in the *in vivo* cortices by scRNA-seq (B) The congruent relationship of *in vitro* and *in vivo* cell types. The unmatched *in vivo* cell types are not shown. (C) Representative images showing the appearance of the deeper-layer marker CTIP2 (red) in converted neurons (GFP^+^DCX^+^) at 5 DPI in ND1 group. Scale bar = 50 μm. (D) The neuronal trajectories captured *in vitro* and *in vivo*. Relevant cell types are shown. (E) Comparison of the two trajectories’ genes. GO terms of the three group of genes are shown on the right. (F) Consistency of expression patterns of the 358 genes between *in vivo* and *in vitro* trajectories. (G) The expression level of endogenous *Ascl1* in *in vivo* and *in vitro* cells.

We further investigated whether the regulatory gene modules identified *in vitro* was also involved in *in vivo* neurogenesis. We found that the turquoise module, associated with astrocyte differentiation (Figure S3D-S3F), was highly active in astrocytes *in vivo* (Figure S5F), while the blue module, related to neurogenesis, was highly active in IPCs and migrating neurons (Figure S3D-S3F and S5F). The yellow module, associated with immature astrocytes, was highly active in dividing RG cells and IPCs. To examine this in more details, we performed pseudotime analysis of the *in vivo* cells and identified a neurogenesis trajectory originating from RG cells (Figure S5G and S5H). When extracting the neurogenesis trajectory genes from the *in vitro* and *in vivo* datasets (Figure 4D, Table S1), we found 358 genes were shared between the two systems, accounting for 64% of the pseudotime genes observed *in vivo* (Figure 4E, Table S1). Most of the shared genes were categorized into biological pathways related to neuronal development and differentiation (Figure 4E). Among these 358 shared trajectory genes, 67% exhibited the same expression pattern along the pseudotime trajectory (Figure 4F and S5I). These findings suggest a high degree of similarity between the *in vitro* neuronal reprogramming and *in vivo* neurogenesis. Meanwhile, the *in vitro* reprogramming also exhibited unique regulatory modes. For example, there were 1821 genes involved in response to innate immune and oxidative stress, which were absent in *in vivo* neurogenesis (Figure 4E), suggesting that those genes were likely induced by viral infection and cell fate switch from astrocytes to neurons. Notably, *Ascl1*, which is absent in *in vivo* cortical excitatory neurogenesis, was transiently upregulated in the early reprogrammed cells (Figure 4G). The temporal increase of *Ascl1* has also been reported in the conversion of glial cells to neuroblasts *in vivo*, suggesting some common regulatory pathways during AtN conversion (Magnusson et al., 2020; Niu et al., 2013; Zhang et al., 2022). In summary, the transcriptional regulation governing ND1-induced neuronal reprogramming of astrocytes is highly similar, but not completely identical, to that observed in *in vivo* cortical deeper-layer neurogenesis.

### High correlation between cell proliferation and cell conversion

A previous study found that cell division was not required for neuronal reprogramming induced by Neurogenin-2 (NGN2) (Heinrich et al., 2010). In the paradigm of cortical neuron development, ND1 acts downstream of NGN2 and participates in terminal neuronal differentiation (Hevner et al., 2006). Therefore, we hypothesized that ND1 might rapidly drive immature astrocytes to exit the cell cycle. However, we unexpectedly observed a higher rate of proliferation in Neu_1 cells compared to Ast_1 cells (Figure 5A), suggesting that cell division may be coupled with ND1-induced trans-differentiation. To investigate this hypothesis, we used time-lapse live imaging to trace cell fate and observed cell division during neuronal reprogramming (Figure 5B). Consistent with the sequencing results, over half of the start cells in the GFP and ND1 groups underwent at least one round of division (Figure 5C and S6A), indicating that the majority of start cells were mitotic. Additionally, we observed that higher proportion of cells underwent cell division in the ND1 group compared with the GFP group (ND1 group: 83% vs. GFP group: 69%, Figure 5C).

**Figure 5.**
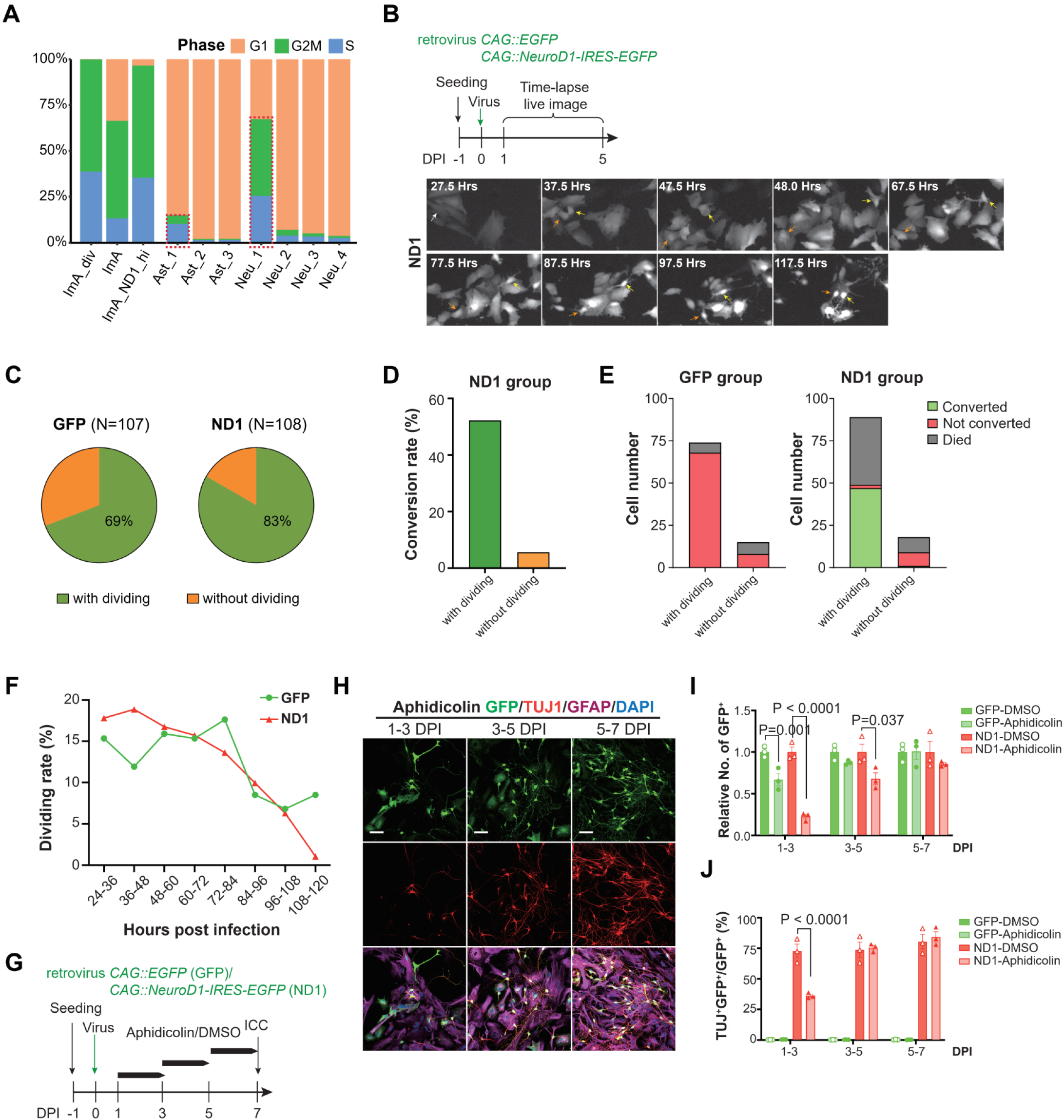
Cell dividing is necessary for successful conversion from ImA to neuron. (A) Ratios of cell cycling states in each cell type. Ast_1 and Neu_1 indicate the two initial differentiation states from ImA. The red dotted boxes indicate more cycling cells in Neu_1 than in Ast_1. (B) Captured pictures of live cell imaging tracing the cell diving during neuronal reprogramming. The arrows indicate the traced cells. (C) Ratios of dividing cells for all the cells we traced in each group. (D) The conversion rates of cell with and without dividing in ND1 group. (E) Number of cells with different final fate for those went through dividing and non-dividing in each group. Two cells without dividing died after conversion were defined as “died” fate. (F) The dividing rate (number of dividing events within the indicated time window/the total number of dividing events) of each group at different time windows after virus infection. (G) Schematic of inhibiting cell cycle experiment. (H-J) Representative images (H) and quantitation show that Aphidicolin abolished neuronal conversion (J) and induced cell death (I) when added at early stage of the reprogramming. Scale bar = 50 μm. One-way ANOVA with Tukey’s multiple comparison test.

When examining neuronal reprogramming based on morphological changes (see Experimental Procedure), we observed that the majority of converted neurons were derived from cells undergoing cell dividing (Figure 5D, SupMoive1). Furthermore, almost all infected cells in ND1 group without cell division were either died or not converted at the end of tracing (Figure 5E). we identified a robust time window for cell division, with the peak occurring within the first 48 hours after transduction. During this period, a higher number of cell division events was observed in the ND1 group (Figure 5F). Subsequently, cell division decreased in both groups, with a sharper decline in the ND1 group. Among the successfully converted cells, over 80% of them underwent one or two divisions, resulting in the generation of 1-2 neurons in most cases (Figure S6B and S6C). The occurrence of cell division in the early stage of neuronal reprogramming was further supported by KI67 immunostaining (Figure S6D-S6G). More KI67^+^ signals were observed in cells transduced with ND1 compared with those transduced with GFP at 2 and 3 DPI (Figure S6E). Taken together, these findings indicate that cell dividing is highly corelated to ND1-induced neuronal reprogramming in this *in vitro* conversion system.

To illuminate the importance of cell dividing for the ND1 induced neuronal reprogramming, we intercepted cell cycle using Aphidicolin, a reversible DNA polymerase inhibitor intercepting cell cycle in S phase (Figure 5G). When added to the robust cell dividing time window (1-3 DPI), Aphidicolin inhibited cell proliferation and significantly inhibited neuronal reprogramming (Figure 5H-J). Whereas, when Aphidicolin was added at 5-7 DPI, the conversion rate was not impaired (Figure 5H-J and S6H). Therefore, cell dividing might be an important factor for ND1-induced neuronal reprogramming from immature astrocytes in cell culture.

### The cellular and molecular regulations of ASCL1-induced neuronal reprogramming

Besides ND1, ASCL1 is another popular TF to induce conversion from astrocytes to neurons, both *in vitro* and *in vivo* (Liu et al., 2015). To investigate the common and specific features of neuronal reprogramming induced by these diverse TFs, we also established *in vitro* AtN conversion system using ASCL1 (Figure 6A and S7) and performed single-cell multiomics analysis (Figure S8A). We observed that ASCL1-induced neuronal reprogramming was slower and less efficient compared with ND1 (Figure 6B and S7). Even at 30 DPI, the neurons reprogrammed by ASCL1 remained in an immature state (Figure S7D, S7E). Furthermore, the subtypes of ASCL1-reprogrammed neurons were predominantly inhibitory neurons (Figure S7F). By analyzing the scRNA-seq data, we identified four distinct cell states along the ASCL1-induced neuronal reprogramming trajectory: ImA, Neu_pre, Neu_a, and Neu_b (Figure 6C-6E). Neu_pre cells exhibited co-expression of ImA genes and proneuronal genes such as *Hes6* and *Sox4* (Table S1). As the reprogramming progressed, ImA genes were down-regulated in Neu_a and Neu_b cells, while Neu_b cells showed elevated expression of genes associated with neuronal development and differentiation, including *Tubb3* and *Stmn3* (Figure S8B-S8C, Table S1). Interestingly, the expression of endogenous Ascl1 was less activated (Figure S8C). When mapping the ASCL1-induced cell clusters to *in vivo* cell types, we found that the converted cells were assigned to Meis2-positive inhibitory neurons (Figure 6F), representing a distinct cell fate compared to the ND1-converted neurons.

**Figure 6.**
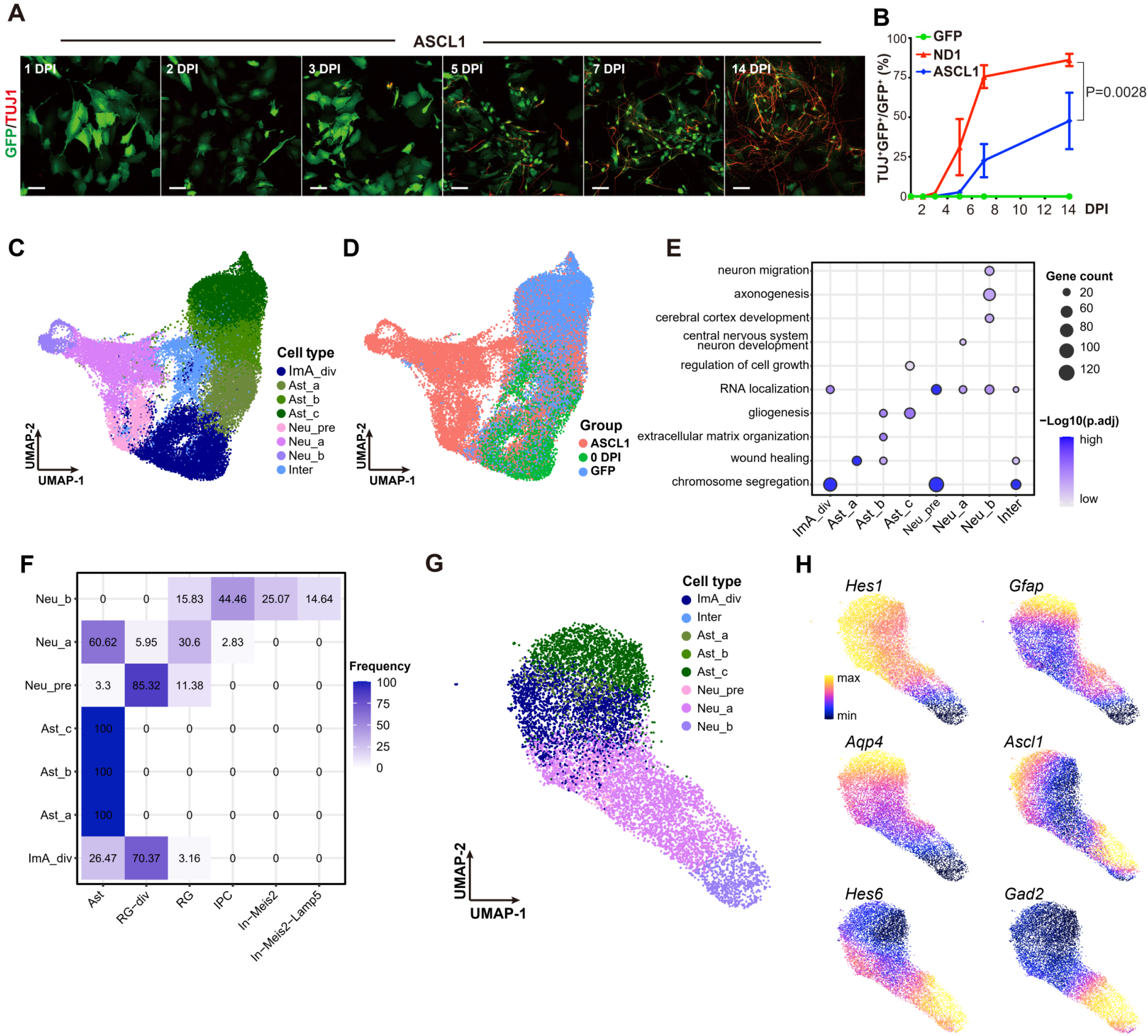
Ascl1-induced *in vitro* neuronal reprogramming. (A) Representative images revealing the gradual morphology change and expression of neuronal marker TUJ1(red) in the Ascl1 group. Scale bar = 50 μm. (B) Conversion rates of cells transfected with ND1 and ASCL1 along time. Two-way ANOVA with Sidak’s multiple comparisons test. (C) UMAP showing the cell types identified using scRNA-seq. (D) The group information of the cells. (E) Representative GO terms of cell type DEGs. (F) The congruent relationship of ASCL1-induced cell types and *in vivo* cell types. The unmatched *in vivo* cell types are not shown. (G) UMAP showing the cell types identified using scATAC-seq. (H) Gene scores of cell type markers.

Then scATAC-seq analysis was performed to investigate the epigenetic changes underlying ASCL1-induced astrocyte fate transition (Figure 6G). Similar to the ND1-induced chromatin changes (Figure S4F), the accessibility landscape revealed distinct patterns for astrocyte-specific genes (*Gfap* and *Aqp4)* in all astrocyte subtypes, while proneuronal genes (such as *Hes6*) were accessible only in neurons (Figure S6H). A number of TFs in ASCL-1 induced AtN reprogramming were predicted, most of which were same with those in ND1 group (Figure S8D).

### Direct comparison of ND1 and ASCL1-induced neuronal reprogramming

Direct comparison of cells from the ND1 and ASCL1 induced neuronal lineages revealed distinct trajectories and final cell fates (Figure 7A, 7C). Pseudotime analysis showed that neurons reprogrammed by ND1 were more mature compared to those reprogrammed by ASCL1 up to 5 DPI (Figure 7B), which was consistent with the observed differences in electrophysiological properties (Figure S7E). Moreover, a higher proportion of proliferative cells was observed in the ASCL1 group even at relatively late stages (5 and 7 DPI, Figure S9A-S9D). Both common and specific genes were identified between ND1 and ASCL1-induced conversions. Genes including *Hes6, Atoh8,* and *Sox11*, known as common core factors involved in AtN reprogramming (Masserdotti et al., 2015), were up-regulated in both groups. However, ND1 induced the expression of these common genes at higher levels compared with ASCL1 (Figure 7B). Other reported factors such as *Neurod4* and *Insm1* were specifically expressed in the ND1 group (Figure 7B). The inefficient increase of these common factors critical for neuronal reprogramming may contribute to the slower and less efficient reprogramming induced by ASCL1. Furthermore, among *Klf10, Chd7,* and *Myt1,* previously reported as critical factors for ASCL1-induced neuronal reprogramming, *Cdh7* and *Myt1* were upregulated in both ND1 and ASCL1-induced conversion, whereas *Klf10* was only transiently upregulated in ASCL1-induced conversion. (Rao et al., 2021) (Figure 7B). The expression of neuronal subtype markers, including *Neurod2* and *Gad2*, was specifically increased in the later stages of reprogramming in the ND1 and ASCL1 groups, respectively (Figure 7B). Unlike ND1, exogenous ASCL1 only slightly promoted the expression of endogenous *Ascl1* at the early stage, and this transient expression of endogenous *Ascl1* was also observed in the ND1 group. However, endogenous *Neurod1* was not promoted by ASCL1. Overall, these findings highlight both common and distinct features of neuronal reprogramming induced by ND1 and ASCL1, providing insights into the different transcriptional programs and cellular outcomes mediated by these TFs.

**Figure 7.**
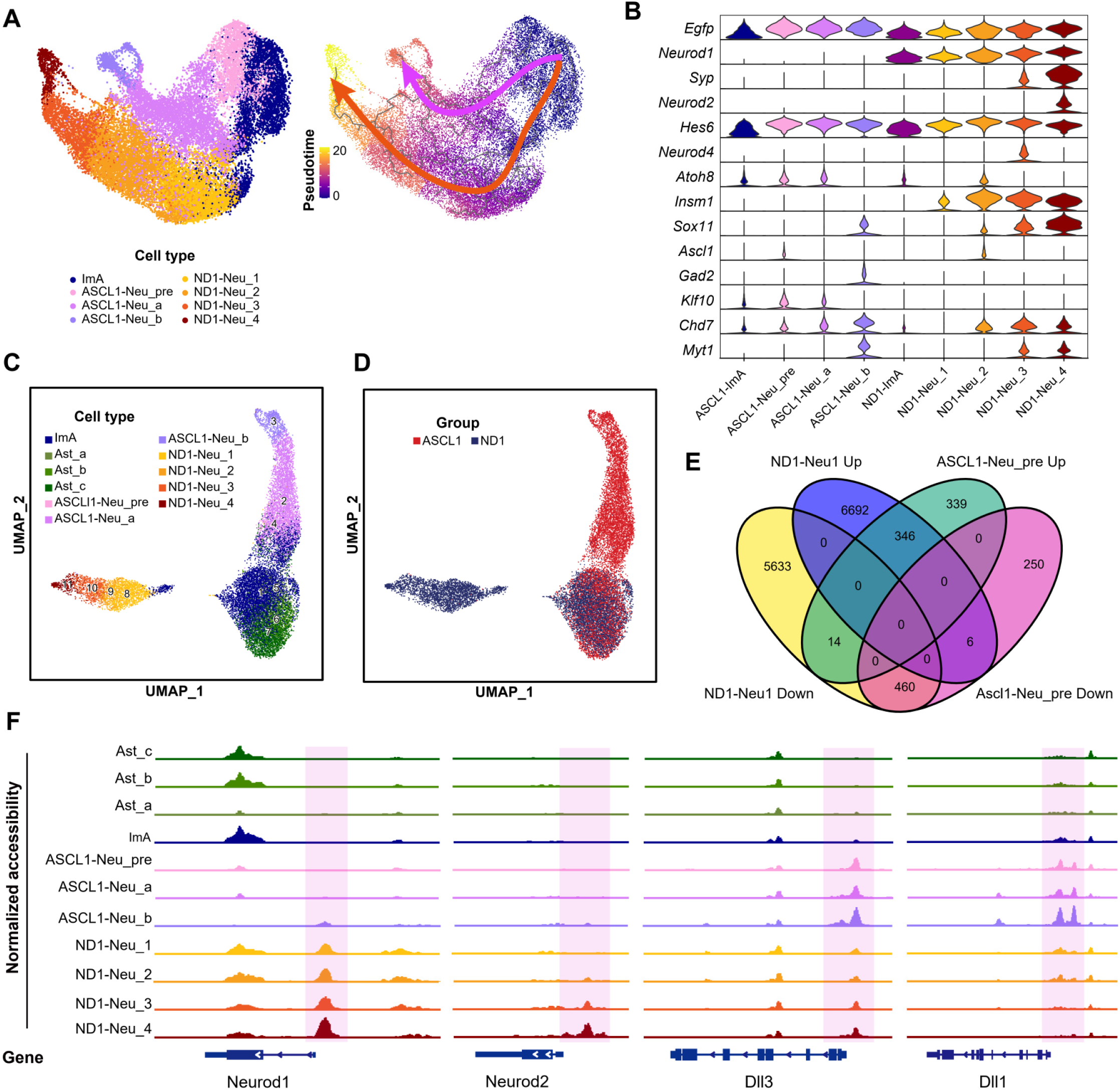
Differences between ND1 and ASCL1-induced astrocyte to neuron reprogramming. (A) UMAP of scRNA-seq data from both ND1 and ASCL1 reprogrammed cells (left). Pseudotime of the cells reveals two independent trajectories (right). (B) Violin plot showing the expression levels of important genes in ND1 and ASCL1 relevant cell types. (C) UMAP of scATAC-seq data including cells from both ND1 and ASCL1 groups. (D) Group information of the cells. (E) Comparison of DARs between ND1-Neu_1 and ImA with those between ASCL1-Neu_pre and ImA. (F) The genome browser view showing the chromatin accessibility signal in each cell type.

According to the scATAc-seq results, both ND1 and ASCL1 were able to induce changes in the chromatin accessibility of ImAs, but ND1 induced more pronounced changes compared with ASCL1 as reflected by the distance between reprogrammed neurons and initial astrocytes (Figure 7C, 7D). The number of DARs between ND1-induced Neu_1 cells and ImAs was over ten times higher than that between ASCL1-induced Neu_pre cells and ImAs (Figure 7E). More than half of the ASCL1-induced DARs were also observed in the ND1 group. For example, *Dll3*, a known target of ASCL1, exhibited open chromatin accessibility in both groups of induced neural cells (Figure 7F). While common putative regulatory TFs were identified (Figure S4H, S8D), there were also TF-specific DARs. *Neurod1* and *Neurod2* peaks were exclusively detected in ND1-induced neurons, whereas *Dll1*, a known target of ASCL1, displayed increased accessibility in ASCL1-induced neural cells (Figure 7F). GO analysis of the DAR associate genes revealed distinct functional enrichment patterns. ND1 initially up-regulated genes involved in telencephalon development and neuron differentiation, while suppressing genes associated with cell migration (Figure S9E and S9F). On the other hand, ASCL1 initially up-regulated genes related to membrane assembly and suppressed genes involved in the ERK1/ERK2 cascade and nervous system development (Figure S9G and S9H). These results suggest that ND1 more efficiently promotes chromatin remodeling favoring neuronal reprogramming compared with ASCL1. ND1 directly promotes neuronal differentiation, whereas ASCL1 initially prevents immature astrocyte from undergoing maturation. Overall, these findings highlight the differences in the reprogramming dynamics, maturation, and neuronal subtype specification between ND1 and ASCL1, underscoring the importance of specific TFs in driving neuronal reprogramming towards different cellular outcomes.

## Discussion

The present study used multiomics combined with time-lapse live image to depict how astrocytes isolated from postnatal rat cortex were converted into functional neurons by ND1 and unravel the underline transcriptomic and epigenetic mechanisms. we discovered an interesting intermediate state co-expressing astrocytic and neuronal genes at the initial stage of neuronal reprogramming. The features of astrocytes were then gradually shut down while the neurogenic program continued. Importantly, the subsequent progress of neuronal conversion recapitulated the transcriptional program of cortical neurogensis. Cell dividing is highly co-related to this indirect AtN conversion. Finally, the integrative multiomic analyses uncovered distinct epigenetic and transcriptional programs between the ND1 and ASCL1-induced neuronal reprogramming. Therefore, this work sheds light on the underline mechanisms of ND1-induced immature astrocyte to neuron conversion.

Previous studies have reported direct and indirect paths of neuronal reprogramming. The direct neuronal reprogramming bypasses the NSC stage, while in the indirect path, astrocytes undergo dedifferentiation into transiently amplifying cells that express ASCL1, which then further differentiate into neuroblasts (Heinrich et al., 2010; Niu et al., 2015). In our ND1-induced AtN conversion, we observed a different pattern to the previous two paths. Upon the stimulation of ND1, the astrocytes entered an intermediate state with simultaneous activation of both astrocytic and neuronal genes. The presence of such intermediate state might be partly due to the characteristics of the initial cells. The cultured astrocytes were isolated from postnatal brains. Although they have been sub-cultured for several passages, these cultured astrocytes still remained as immature astrocytes with high proliferation ability (Lattke et al., 2021). In the absence of pro-neuronal TFs, they would gradually mature under prolonged culture, as revealed by the astrocyte branch in our pseudotime analysis. This default program of astrocyte maturation might not be immediately shut down by the exogenous ND1 and other downstream effectors could help to repress the astrocyte genes.

Multiomics analyses revealed larger number of pro-neuronal genes directly upregulated by ND1, suggesting that ND1 might predominantly play a role as activator of neurogenic program. As the pro-neuronal genes (e.g. *Hes6*) accumulated, the intermediate cells gradually shut down astrocytic genes and adopted a neurogenic program. An interesting observation is a transient activation of *Ascl1* during this stage. Activation of *Ascl1* is not involved in the development of cortical pyramidal neurons, but it has been reported transiently elevated in the indirect parenchymal AtN conversion (Magnusson et al., 2020; Zhang et al., 2022). So there might be some common regulations among the intermediate states induced by different methods. Once neurogenic program initiated, this ND1-induced AtN conversion recapitulates the transcriptional progress of cortical deeper-layer neuron genesis, passing through RGs, IPCs, and migrating neurons. Probably because of this high similarity between these two programs, ND1-converted neurons shared the characteristics of primary cortical deeper-layer neurons both in gene expression and electrophyisiological features.

Another interesting finding in this study is that cell dividing is highly correlated to successful conversion. The time-lapse live image results show that over 99% converted neurons derived from cells underwent cell dividing, leading us to investigate whether cell dividing might have a relationship with cell conversion. Indeed, we discovered that during the conversion, cells undergoing cell dividing were more likely converted into neurons than those without cell dividing, suggesting that cell dividing might be a critical event to the complete AtN conversion in this study. This is consistent with previous studies reporting that dividing astrocytes infected by ND1 retroviruses are relatively easier converted into neurons(Guo et al., 2014) compared to postmitotic resting astrocytes infected by AAV (Brulet et al., 2017). It is of great interest and scientific merit to investigate the dynamic progress and gene regulations in neuronal reprogramming of postmitotic astrocytes in the future.

Regarding ASCL1-induced neuronal conversion, we also found that it shared partial features with the *in vivo* neurogenic program of forebrain interneurons. But compared with ND1, the conversion induced by ASCL1 was lower and inefficient. Through comparison of the transcriptome and epigenome between the two TF infected cells, we discovered that the reported critical common neurogenic factors (*Hes6*, *Atoh8, Sox11*, *Neurod4*, and *Insm1*) (Masserdotti et al., 2015) were less or not elevated in ASCL1 group. In summary, our single-cell multiomic analyses discover fundamental mechanisms underlying the conversion from an astrocyte to a neuron after overexpressing a single neuronal TF, NEUROD1. The distinct transcriptomic pathways between ND1- and ASCL1-induced AtN conversion also provide a potential framework for future *in vivo* applications in neuroregeneration and neural repair.

## Acknowledgements

We thank Prof. Minwei Min (Guangzhou National Laboratory) for the help in data collection of live cell imaging. This work is supported by grants from the National Key Research and Development Program of China (2020YFA0112201 to X.F.), the Key Research and Development Program of Guangzhou (202206060002 to X.F., W.L. and G.C.), and the Guangdong Provincial Pearl River Talents Program (2021QN02Y747 to X.F.).

## Author Contributions

X.F., W.L. and G.C. conceived the ideas. X.F. and W.L. supervised the project. W.L., and K.L. established the *in vitro* neuronal reprogramming platform, collected the data and finished the analysis except sequencing experiments, with the assistant of D.S.. D.S. performed all the single-cell experiments and X.Li. did the bioinformatics analyses. Q.P. did the RT-qPCR validation experiments. J.Z. collected the data of electrophysiology. X.Luo constructed the Retroviral vectors. X.F., W.L., D.S. wrote the manuscript, and all authors discussed the data. X.F., G.C., W.L. proofread the manuscript.

## Data and code availability

The raw data used in this paper have been deposited to the Genome Sequence Archive in National Genomics Data Center with the accession number PRJCA017801. No customized code is used in this study and code used throughout this study is available upon reasonable request from the authors.

## EXPERIMENTAL PROCEDURES

### Primary cell culture

Astrocytes were cultured as previously described with several modifications [1]. Briefly, the cortices from postnatal day 2-3 Sprague Dawley rat were dissected and dissociated with 0.15% trypsin-EDTA (a mixture of 0.05% Trypsin-EDTA:0.25% Trypsin-EDTA at 1:1 volume ratio) for 15 min. The cell suspension was then seeded in non-coated flasks for expansion with the medium containing DMEM/F12 (supplemented with 4.5 g/L glucose, 2 mM L-glutamine), 10% fetal bovine serum (FBS, Australia origin) and 1% penicillin/streptomycin in a 5% CO2 and 37°C incubator (All reagents provided by Thermo Fisher scientific, Grand Islands, NY). After 7-9 days, cell confluence reached ∼90%. Non-astrocytic cells, including microglia, neurons, oligodendrocytes and their progenitors, were vigorously shaken off and the attached cells were reseeded in astrocyte maintenance medium containing DMEM/F12 (supplemented with 3.5 mM glucose, 2 mM L-glutamine), 2% B27, 10% FBS and 1% penicillin/streptomycin. The astrocyte culture was passaged 4 times to eliminate the neural stem cells before subsequent experiments.

Primary neurons were isolated from cortices of Sprague Dawley rat E16.5 pups as described previously [2]. Briefly, the cortices were dissected in ice cold artificial CSF (pre-bubbled with 95%O2::5%CO2), cut into fine pieces and incubated with 7.5 units/ml papain solution containing L-cysteine (1 mM), EDTA (0.5 mM) and DNase I (150 units/ml), dissolved in EBSS (equilibrated with 95%O2::5%CO2) at 34 °C for 30 min. Then, the tissue was triturated with 200 μL pipet tips and filtered through a 70 µm cell strainer (BD, Franklin Lakes, NJ). Cell pellet was collected by centrifuged at 300 g for 5 min and then resuspended in 3 mL Ovomucoid protease inhibitor with FBS (Inhibitor-BSA Vial [LK003182], Worthington, Lakewood, NJ). To remove the debris, the cell suspension was further centrifuged at 70 x g, 6min. The cell pellets were resuspended in the medium containing DMEM/F12 (supplemented with 3.5mM glucose, 2 mM L-glutamine), 2% B27, 1% FBS and 1% penicillin/streptomycin and seeded at the density of 15,000-20,000 cells/12mm poly-D-Lysine coated coverslip. 5 days later, when the neurites appeared, 200 μg/mL L-ascorbic acid (Merck/Sigma-Aldrich), 1 μM cyclicadenosine monophosphate (Merck/Sigma-Aldrich), 1 μg/mL laminin (Merck/Sigma-Aldrich), 20 ng/mL brain-derived neurotrophic factor (Peprotech, Rocky Hill, NJ, USA), 10 ng/mL neurotrophin-3 (Peprotech), and 10 ng/mL insulin-like growth factor (Peprotech) were added and the medium was changed every 2-3 days.

All animal procedures have been approved by Jinan University Institutional Animal Care and Use Committee (Approval No. IACUC-20180330-06).

### Retrovirus production

Retroviral vectors, CAG::GFP (GFP retrovirus), and CAG::NeuroD1-IRES-GFP (NEUROD1-GFP retrovirus) were obtained from the previous study [3]. Retroviral vector CAG::Ascl1-IRES-GFP (ASCL1-GFP retrovirus) was constructed by replacing NeuroD1 with the open reading frame of mouse Ascl1. All viruses were packaged and concentrated as previously described[3]. The titer of viral particles was about 1 × 10^8^ transfer units/mL, which was determined after transduction of HEK293T cells.

### *In vitro* AtN conversion

Primary astrocytes at passage 5 were seeded at the density of 10,000–12,000 cells on poly-D-lysine (Merk/Sigma-Aldrich)-coated glass coverslips (12 mm in diameter, Glaswarenfabrik Karl Hecht GmbH &Co., Sondheim, Germany) in astrocyte maintenance medium. After culturing for 24 hours, when the cell confluence reached 70-80%, the retrovirus was added at 5-10 MOI. Next day, the medium was switched to conversion medium containing DMEM/F12 supplemented with 3.5 mM glucose, 2 mM L-glutamine, 2% B27, 1% FBS, and 1% penicillin/streptomycin. Five days post-infection when neurite-like processes appeared, 200 μg/mL L-ascorbic acid (Merck/Sigma-Aldrich), 1 μM cyclicadenosine monophosphate (Merck/Sigma-Aldrich), 1 μg/mL laminin (Merck/Sigma-Aldrich), 20 ng/mL brain-derived neurotrophic factor (Peprotech, Rocky Hill, NJ, USA), 10 ng/mL neurotrophin-3 (Peprotech), and 10 ng/mL insulin-like growth factor (Peprotech) were added. During the conversion, half the medium was changed every other day. To inhibit the cell proliferation, Aphidicolin (5 μM, Abcam) was added into the conversion medium at indicated time points.

To investigate the mode of the cell death during AtN transdifferentitaion, Z-VAD-FMK (20 μM, Selleckchem), Liproxstatin-1 (200 nM, Selleckchem), or Necrostatin-2 (10 μM, Selleckchem) was added to the conversion medium from 2 DPI to 5 DPI.

### Reverse-transcription and real-time quantitative PCR

Cells were harvested to extract RNA using ReliaPrep™ RNA Cell Miniprep System (Promega). CDNA was reverse-transcribed using PrimeScript™ RT reagent Kit (Takara). Real-time quantitative PCR was performed using QuantiNova® SYBR® Green RT-PCR Kit (Qiagen) according to the manuals on CFX Real-Time qPCR system (BioRad).

### Immunofluorescence

Cells were fixed with 4% paraformaldehyde for 15 minutes and then incubated in 0.01% Triton X-100 in PBS for 10 minutes at room temperature. After three washes with PBS, 3% bovine serum albumin (BSA, Merck/Sigma-Aldrich) in PBS was added as blocking buffer and incubated for 1 hour. Then cells were incubated with the indicated primary antibodies diluted in 1% BSA at 4°C overnight. 0.2% PBST (Tween-20 in PBS) was used to wash away the unbound antibodies. Next, 1:1000 diluted secondary antibodies were added and incubated for 1 hour at room temperature. Finally, after washed by 0.2% PBST three times, 0.5 μg/mL DAPI (4,6-diamidino-2-phenylindole, F. Hoffmann-La Roche, Natley, NJ, USA) was added to counterstain the nuclei. The coverslips were mounted on glass slides using anti-fading mounting medium (DAKO, Carpinteria, CA, USA).

Brains were collected and fixed with 4% paraformaldehyde at 4°C for 2-6 hour. After fixation, the tissues were washed in cold PBS three times and transferred to 30% sucrose at 4°C until sank. The tissues were then embedded with O.C.T. (Tissue-Tek®, Torrance, CA) and cryosection at 15 μm thickness. The sections were washed 3 times with PBS before subjected to antigen retravel (in 95°C citrate buffer for 10 min). Sections were incubated in blocking buffer (5% normal donkey serum, 3% BSA, and 0.2% PBST) at room temperature for 1 hour and then incubated with primary antibodies for 24-48 hours at 4°C. Thereafter, brain sections were rinsed with 0.2% PBST and incubated with corresponding secondary antibodies and DAPI for 2 h at room temperature, followed by an extensive wash with 0.2% PBST. Finally, the immunofluorescence stained brain sections were mounted with mounting medium (VECTASHIELD®, VECTOR Laboratories, Burlingame, CA, USA) and sealed with nail polish.

Images were collected with a fluorescence microscope (Axio Imager Z2, Zeiss) for quantification and with a confocal microscope (LSM880, Zeiss) for representative image display.

### Electrophysiological recording

Whole-cell recordings were performed on transdifferentiated neurons at 30 DPI or primary neurons at 30 DIV (days *in vitro*) using Multiclamp 700A patch-clamp amplifier (Molecular Devices, Palo Alto, CA) as described before[4], and the chamber was constantly perfused with a bath solution consisting of 126 mM NaCl, 2.5 mM KCl, 1.25 mM NaH2PO4, 26 mM NaHCO3, 2 mM MgCl2, 2 mM CaCl2 and 10 mM glucose. The pH of bath solution was adjusted to 7.3 with NaOH, and osmolarity was at 310–320 mOsm/L (all reagents provided by Merck/Sigma-Aldrich). Patch pipettes were pulled from borosilicate glass (3–10 M) and filled with a pipette solution consisting of 126 mM K-Gluconate, 4 mM KCl, 10 mM HEPES, 4 mM Mg2ATP, 0.3 mM Na2GTP, 10 mM PO Creatnine (pH 7.3 adjusted with KOH, 290 mOsm/L). For voltage-clamp experiments, the membrane potential was typically held at −70 or −80 mV. Data was acquired using pClamp 10 software (Molecular Devices, Palo Alto, CA), sampled at 20 kHz, and filtered at 3 kHz. Na+ and K+ currents and action potentials were analyzed using pClamp 10 Clampfit software. Spontaneous synaptic events were analyzed using MiniAnalysis software (Synaptosoft, Decator, GA). All experiments were conducted at room temperature.

### Time-lapse live cell image

Time-lapse live cell image of AtN transdifferentiation was performed with Nikon Ti2E Inverted Microscope. Fluorescence images were acquired every 30 min from 1 DPI to 5 DPI using a 20 x phase contrast objective. Videos were assembled using Image J 1.52p (National Institute of Health, USA) software and were played at speed of 6 frames per second.

### Single cell suspension acquisition

For culture cells, the attached cells at the indicated time points were dissociated using 0.05% Trypsin-EDTA to get single cell suspension. The procedure for preparing the single cell suspension from the cortices was similar to that for preparing primary neuron culture. The cell pellets were resuspended in sterile-filtered washing buffer (Dulbecco’s PBS containing sodium pyruvate, streptomycin sulfate, kanamycin monosulfate, glucose and calcium chloride; Sigma-Aldrich, D4031) containing 0.5% BSA.

### NEUROD1 ChIP-seq

GFP^+^ cells were sorted with fluorescence-activated cell sorting (FACS) using BD FACSAria III cell sorter at 2 DPI. 10^6^ cells per test were cross-linked 15min at room temperature by 1% Formaldehyde. Cell pellets were sonicated for 10 min using a Covaris S220 instrument (duty factor 2%; Peak incident power:105w; cycles per bust:200). Chromatin was immunoprecipitated with 2ug of NEUROD1 antibody (Cell Signaling mAb #4373). After DNA purification, ChIP-seq libraries were constructed by KAPA HyperPrep Kit (KK8504).

### ScRNA-seq and scATAC-seq library preparation and sequencing

For scRNA-seq, cells were barcoded through the Singleron Matrix instrument using the GEXSCOPE Single Cell RNA Library Kit contain GEXSCOPE microchip, barcoding beads, and reagents for transcriptome amplification and library construction (Singleron Biotechnologies, 4180012). The sequencing libraries were prepared according to the manufacturer’s instructions and sequenced on Nova6000 of Illumina with PE150 reads. For scATAC-seq, nuclei were isolated according to 10x genomics protocol CG000169 (Demonstrated Protocol Nuclei Isolation ATAC Sequencing Rev E). ScATAC–seq libraries were generated using the Chromium Single Cell ATAC V1 Library & Gel Bead Kit. All libraries were sequenced using MGI2000 with PE100 reads.

### ScRNA-seq data analysis

Raw reads were processed to generate gene expression profiles using CeleScope v1.6.0 pipeline (https://github.com/singleron-RD/CeleScope) with default parameters. Briefly, Barcodes and UMIs were extracted from R1 reads and corrected. Adapter sequences and poly A tails were trimmed from R2 reads. The clean R2 reads were then aligned to the Rattus norvegicus genome (mRatBN7.2) using STAR (v2.6.1b) [5]. Uniquely mapped reads were assigned to exons with FeatureCounts (v2.0.1) [6]. Successfully assigned reads with the same cell barcode, UMI and gene were grouped together to generate the gene expression matrix for further analysis.

Genes detected in less than 10 cells were removed. DoubletFinder (v2.0.3) [7] was used to filter potential doublets for each sample. Cells were discarded if they met any of the following conditions: 1) expressed less than 1000 genes; 2) detected with more than 10% of mitochondrial genes; 3) contained reads number outside the range of 10^(mean(log10(reads number)) ± 2*sd(log10(reads number))).

After stringent quality control, remained cells were analyzed using Seurat package (v4.0.3) [8]. The filtered count matrix was firstly log normalized using NormalizeData() function. Next, top 2000 highly variable genes were extracted by FindVariableFeatures() function, and scaled to compute principal components through ScaleData() and RunPCA(), respectively. The mutual nearest neighbors (MNN) method was used to alleviate the batch effect. Unsupervised clustering was performed on the scaled and batch corrected data by FindNeighbour() and FindCluster() function using the top 20 PCs. Uniform Manifold Approximation and Projection (UMAP) was employed to visualize the result of clustering. Cellular state labels were assigned to each cluster based on marker genes reported by FindAllMarkers() function, and we manually validated these cell state labels according to previously reported marker genes, such as *Gfap* for astrocytes and *Dcx* for newborn neuron. ClusterProfiler (v3.4.0) [9] was used to characterize each cellular states by Gene Ontology (GO) terms.

### Trajectory analysis

A trajectory graph was constructed using Monocle3 (v3.0) [10] on UMAP coordinates from Seurat. Cells from D0 were selected as root cells. Pseudotime inference was performed using order_cells() function. We also took advantages of Monocle2 [11] to describe the cellular state divergences. To compare the gene expression between two paths, we used branched expression analysis modeling (BEAM) [12] and visualized the results using the plot_genes_branched_heatmap() and plot_genes_branched_pseudotime() function.

### Key transcription factor analysis

To identify the key regulators that drive the differentiation process, we first used ARACNe-AP [13] (v1.0.0) to build transcriptional regulatory networks. In brief, Rattus Norvegicus transcription factors in AnimalTFDB and gene expression matrix from Ast and Neu states, which were described by monocle2, were taken as input to the ARACNe-AP. Then, MARINa algorithms, implemented by R package ssmarina (v1.01) was used to analyze the master regulatory for each differentiation route.

### WGCNA analysis

HdWGCNA (v0.2.17) [14] was used to construct co-expression networks across different cellular states. Briefly, we aggregated similar cells into several small groups by running MetacellsByGroups() function on Seurat object. Soft power threshold was inferred using TestSoftPowers() function. The co-expression network was finally constructed by running ConstrucNetwork() function. The module eigengenes (MEs) were calculated with ModuleEigengenes() function. The hub genes for each module were identified using ModuleConnectivity() and ModuleExprScore() function.

### Mapping *in vitro* cells to *in vivo* references

To annotate *in vitro* query datasets based on the *in vivo* cortical reference data, we first projected the PCA structures of a reference onto the query by running FindTransferAnchors() function. Then, *in vitro* cells were classified based on *in vivo* cell type labels using TransferData() function. To guarantee an accurate annotation, we removed predicted reference cell types in which less than 50 cells in a cell cluster were assigned to. Ggalluvial [15] was used to visualize the prediction results.

### ScATAC-seq data analysis

#### scATAC-seq data processing

Raw sequencing data were converted to fastq format using ‘cellranger-atac mkfastq’ (10x Genomics, v.2.0.0). scATAC-seq reads were aligned to the Rattus norvegicus genome (mRatBN7.2) and quantified using ‘cellranger-atac count’ (10x Genomics, v.2.0.0). Fragment data was loaded into ArchR (v1.0.1) [16] for quality control and downstream analysis. In brief, fragments on Y chromosome and mitochondrial DNA were removed. Cells with less than 1,000 or more than 100,000 fragments were filtered. We additionally identified and discarded potential doublets by using add Doublet Scores () function. To guarantee a high signal-to-noise ratio, cells with a TSS enrichment score less than 4 were also excluded in subsequent analyses.

#### scATAC-seq clustering and dimensionality reduction

To cluster scATAC-seq data and visualize cell embedding in a reduced dimension space, such as UMAP, we first applied iterative latent semantic indexing (LSI) on the top 25,000 accessible 500-bp tiles by running addIterativeLSI() function. Clustering was performed using addClusters() function with ‘resolution’ set as 0.8. An UMAP representation was obtained by running addUMAP() function with ‘minDist’ parameter set to 0.6.

#### Label transfer

To annotate scATAC-seq clusters, we first calculated gene score by running addGeneScoreMatrix() function to estimate gene expression level based on chromatin accessibility data. Then, we implemented canonical correlation analysis (CCA) by performing addGeneIntegrationMatrix() function for a preliminary unconstrained integration of scRNA-seq and scATAC-seq datasets. To further refine the integration results, we determined the most enriched scRNA-seq based cell labels in each of the scATAC-seq clusters, and then performed a second round integration by constraining the scATAC-seq clusters to the most corresponding scRNA-seq based cell types. We validated the label transferred results by known cell type marker genes.

#### scATAC-seq peak identification

Since the extreme sparsity in scATAC-seq dataset, which may hinder the peak identification, we created pseudo-bulk replicates by grouping cells from the same clusters using addGroupCoverages() function. Cluster specific peaks were called using those pseudo-bulk replicates with MACS2 (v2.2.7) [17]with ‘-g’ parameter set to 2.6+10e9. The peaks were visualized using plotBrowserTrack() function.

#### scATAC-seq motif accessibility deviations

We used chromVAR [18] to predict the enrichment of TFs for each cell type. The chromVAR deviation scores were calculated by running addDeviationsMatrix() function. The position weight matrices (PWM) used in the function were obtained from the JASPAR 2018 [19] and JASPAR 2020 database [20].

#### Identification of peak-to-gene links

Peaks were linked to gene based on a correlation approach, which was implemented in ArchR by running add Peak2GeneLinks() function. Briefly, peaks were associated to the TSS of genes within a 250kb genomic distance, and the Pearson correlation was calculated between scATAC-seq and scRNA-seq values. Only peak-to-gene pairs with r > 0.35 were retained.

### ChIP-seq data processing

ChIP-seq reads were firstly quality controlled using Fastp (v0.21) [21]. Then, Bowtie2 (v 2.4.1) [22] was used to align the clean reads to Rattus norvegicus genome with default parameter. Samtools (v1.15.1) [23] was used to convert SAM file into BAM format. BAM file was then sorted and indexed. Duplicate reads in the bam file were identified and removed by Picard (v3.0.0) [24] using MarkDuplicates function. The peaks were called using MACS2 (v2.2.7) with the ‘-g’ parameter set to 2.6e9. BedGraphToBigWig (v2.8) was used to convert Bedgraph (bdg) file into bigwig format, which was then visualized using Itegrative Genomics Viewer (IGV v2.9.4). Homer (v4.11.1) was used for peak annotation.

### Inferring ND1 target genes using ChIP-seq and scRNA-seq data

To obtain confident ND1 target genes, we first selected genes which were bound by NEUROD1 in promoter or distal regions from ChIP-seq dataset. Then, we tested whether the corresponded genes were differentially expressed in neuronal cells defined in scRNA-seq data. Genes with adjusted p-values < 0.05 and |avg_logFC| > 0.5 were considered as potential ND1 targets.

### Integration ND1 and ASCL1 datasets

To integrate scRNA-seq datasets, we first merge the raw gene expression matrices. Then, we used Seurat (v4.0.3) package to normalize and centralize data on this merged dataset. PCA was performed to reduce the dimensionality. Next, MNN was implemented to correct the batch effect on the top PCs by the monocle3 package. UMAP was employed to visualize the merged results.

For the integration of scATAC-seq datasets, fragment data from ND1 and ASCL1 were merged. Then, we took advantage of harmony, which was accomplished by ArchR, to integrate two datasets.

### Quantification and statistical analysis

#### Cell number analysis

For the analysis of cell number changes during AtN transdifferentiation, fluorescent images of live cells were captured by an inverted fluorescence microscope (Zeiss Axio Observer A1) at 100× magnification (929.79 μm × 929.79 μm) at indicated time points. Five randomly chosen fields were calculated and three cell batches were obtained.

#### Immunofluorescence analysis

Quantification of immunostaining was performed by Zeiss ZEN 2.3 software (blue edition, Göttingen, Germany) using images captured at 200× magnification (464.9 μm × 464.9 μm) by a fluorescence microscope (Axio Imager Z2, Zeiss). Parameters for image capturing and post-analysis were adjusted to the same values for each antigen tested. 15–20 random fields per coverslip were chosen and 3–4 coverslips were used per cell batch. Three cell batches isolated in three independent experiments were used.

#### Time-lapse live imaging analysis

Cells were tracked in every frame and proliferation, neuronal conversion or cell death was analyzed for every single cell in each series of images. To quantify the proportion of dead neurons or astrocytes, we counted cells dying as positive events and classified them as neurons or astrocytes according to their morphology described as [23].

#### Real-time qPCR analysis

The data were plotted as means of 3 independent experiment (using 3 batches of cells).

All values were given as mean ± SEM. The data were tested for significance using two-tailed t-test and one, or two-way ANOVA with Tukey’s or Sidak correction for multiple comparisons (Prism 8, GraphPad). *P* < 0.05 was considered statistically significant.

