## Supplementary figures for "Activation of a cortical neurogenesis transcriptional program during NEUROD1-induced astrocyte-to-neuron conversion"

Figure S1

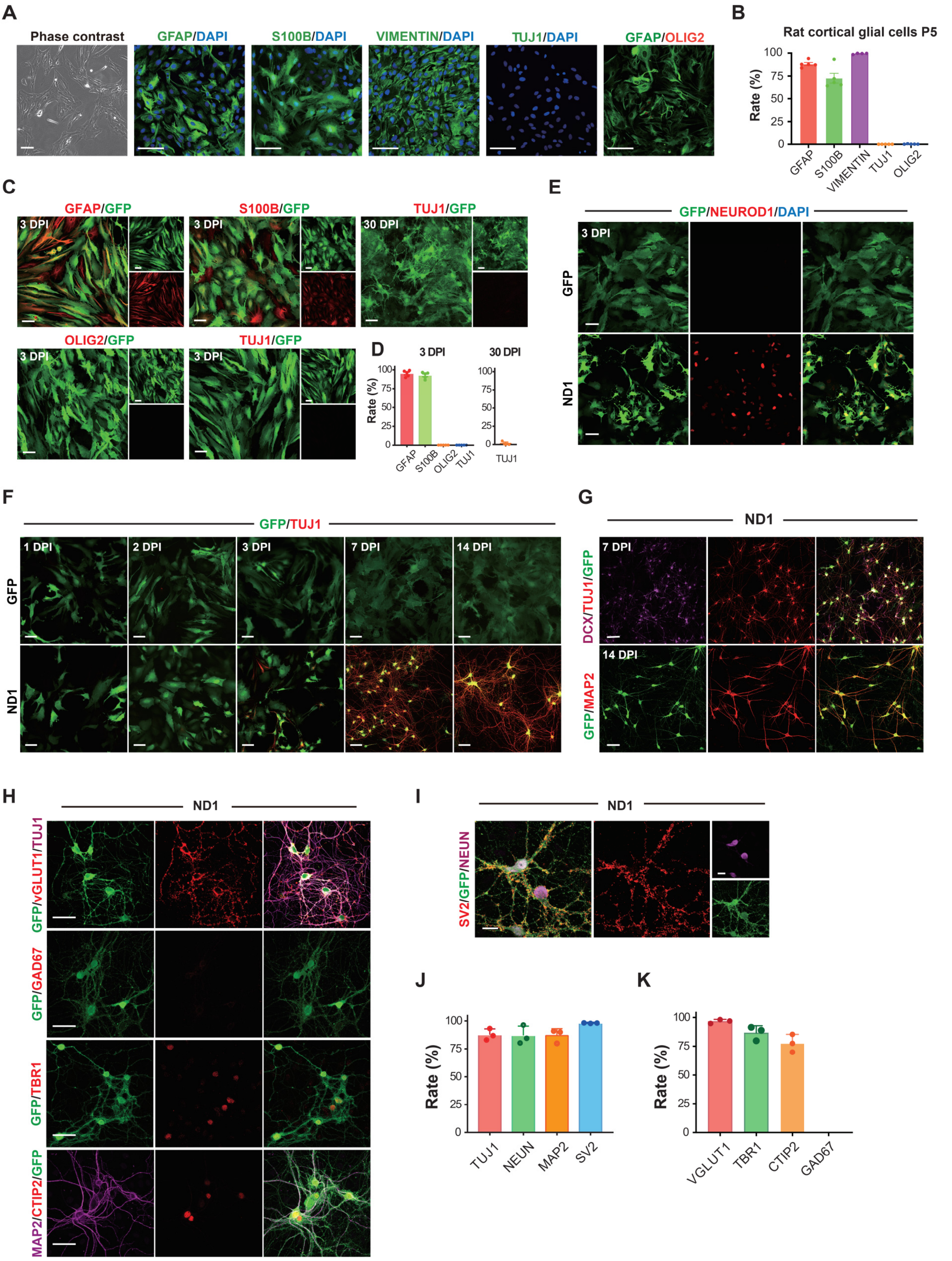
A
Phase contrast
GFAP/DAPI
S100B/DAPI
VIMENTIN/DAPI
TUJ1/DAPI
GFAP/OLIG2
B
Rat cortical glial cells P5
Rate (%)
GFAP
S100B
VIMENTIN
TUJ1
OLIG2
C
GFAP/GFP
S100B/GFP
TUJ1/GFP
OLIG2/GFP
TUJ1/GFP
3 DPI
30 DPI
D
3 DPI
30 DPI
Rate (%)
GFAP
S100B
OLIG2
TUJ1
E
GFP/NEUROD1/DAPI
GFP
ND1
3 DPI
F
GFP/TUJ1
1 DPI
2 DPI
3 DPI
7 DPI
14 DPI
GFP
ND1
G
ND1
DCX/TUJ1/GFP
GFP/MAP2
7 DPI
14 DPI
H
ND1
GFP/vGLUT1/TUJ1
GFP/GAD67
GFP/TBR1
MAP2/CTIP2/GFP
I
ND1
SV2/GFP/NEUN
J
Rate (%)
TUJ1
NEUN
MAP2
SV2
K
Rate (%)
VGLUT1
TBR1
CTIP2
GAD67

Figure S2

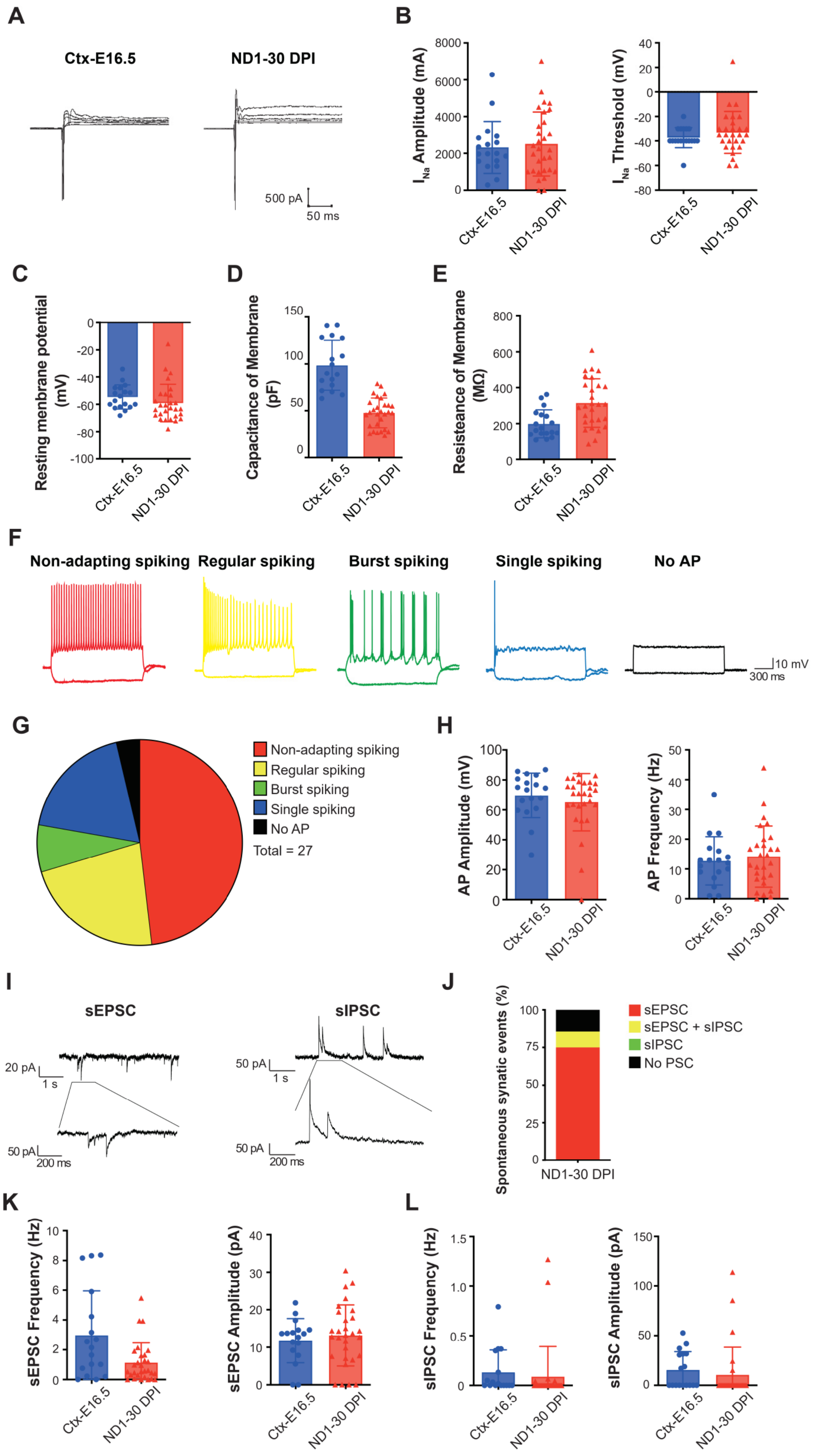
A
Ctx-E16.5
ND1-30 DPI
500 pA
50 ms
B
I_Na Amplitude (mA)
I_Na Threshold (mV)
Ctx-E16.5
ND1-30 DPI
C
Resting menbrane potential (mV)
D
Capacitance of Membrane (pF)
E
Resisteance of Membrane (MΩ)
F
Non-adapting spiking
Regular spiking
Burst spiking
Single spiking
No AP
10 mV
300 ms
G
Non-adapting spiking
Regular spiking
Burst spiking
Single spiking
No AP
Total = 27
H
AP Amplitude (mV)
AP Frequency (Hz)
I
sEPSC
sIPSC
20 pA
1 s
50 pA
200 ms
J
Spontaneous synatic events (%)
sEPSC
sEPSC + sIPSC
sIPSC
No PSC
ND1-30 DPI
K
sEPSC Frequency (Hz)
sEPSC Amplitude (pA)
L
sIPSC Frequency (Hz)
sIPSC Amplitude (pA)

Figure S3

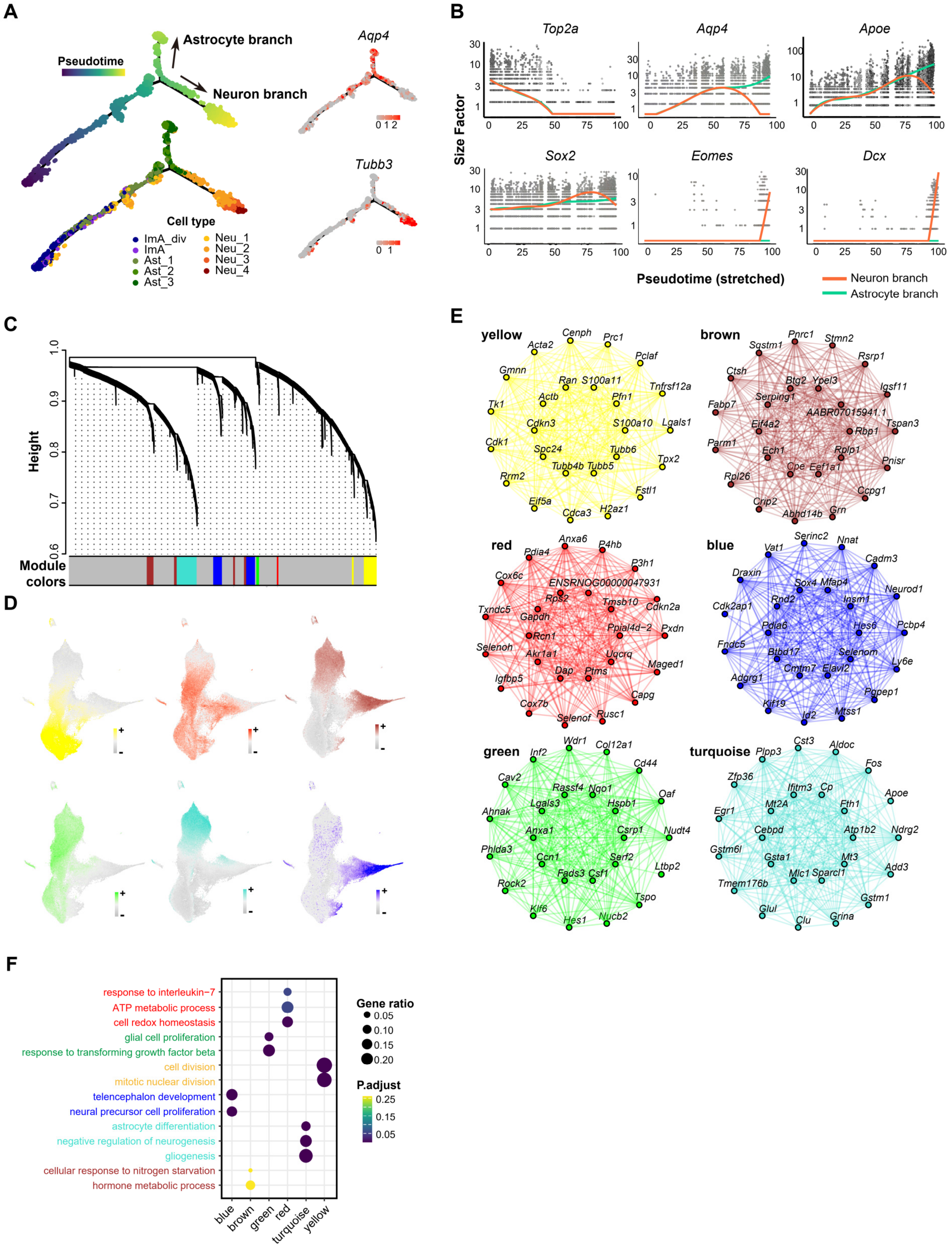

A
Pseudotime
Astrocyte branch
Neuron branch
Aqp4
Tubb3
Cell type
ImA_div
ImA
Ast_1
Ast_2
Ast_3
Neu_1
Neu_2
Neu_3
Neu_4
B
Top2a
Aqp4
Apoe
Sox2
Eomes
Dcx
Size Factor
Pseudotime (stretched)
Neuron branch
Astrocyte branch
C
Height
Module colors
D
E
yellow
brown
red
blue
green
turquoise
F
response to interleukin−7
ATP metabolic process
cell redox homeostasis
glial cell proliferation
response to transforming growth factor beta
cell division
mitotic nuclear division
telencephalon development
neural precursor cell proliferation
astrocyte differentiation
negative regulation of neurogenesis
gliogenesis
cellular response to nitrogen starvation
hormone metabolic process
blue
brown
green
red
turquoise
yellow
Gene ratio
0.05
0.10
0.15
0.20
P.adjust
0.25
0.15
0.05

Figure S4

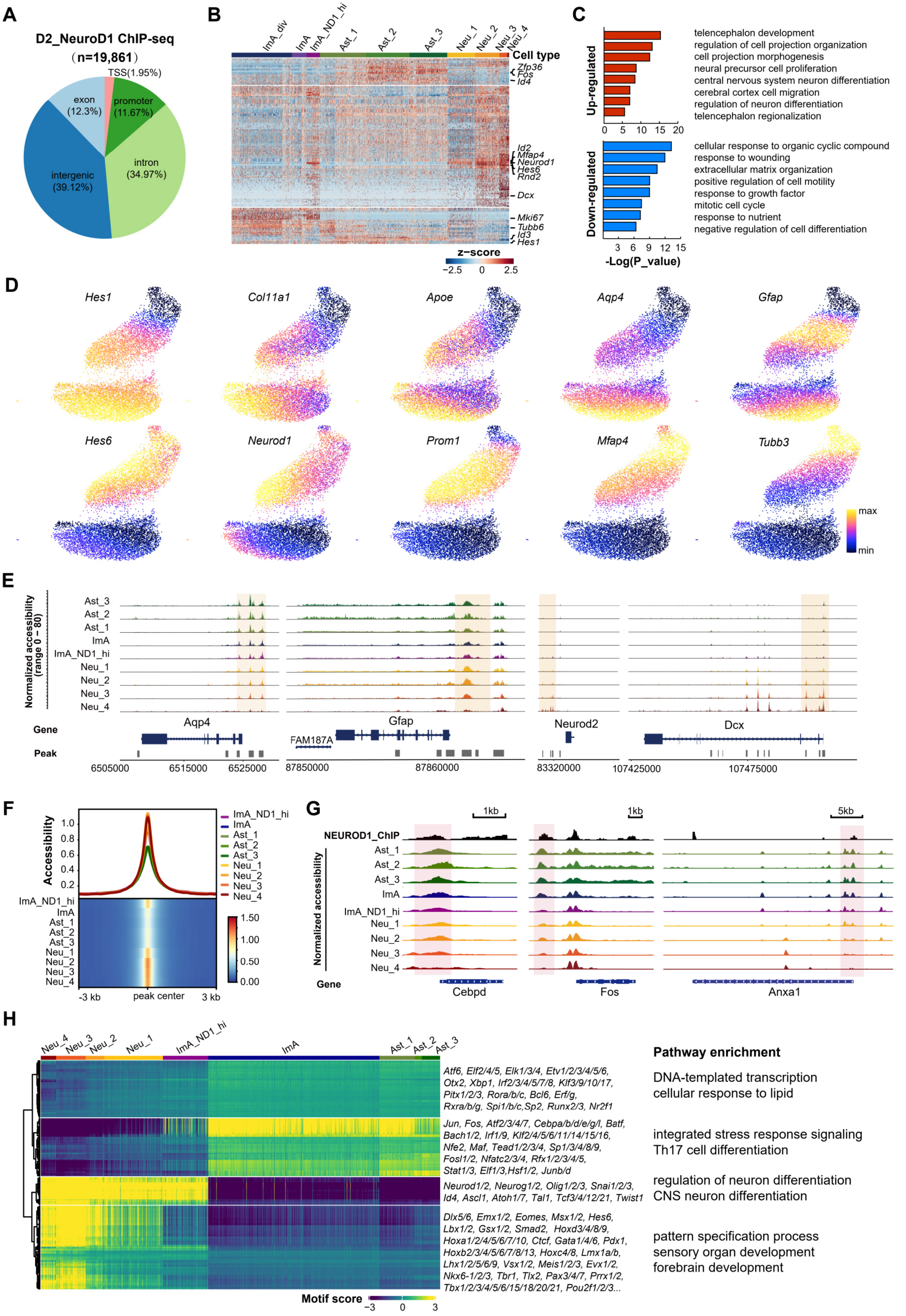
A
D2_NeuroD1 ChIP-seq
(n=19,861)
TSS(1.95%)
exon (12.3%)
promoter (11.67%)
intron (34.97%)
intergenic (39.12%)
B
ImA_div
ImA
ImA_ND1_hi
Ast_1
Ast_2
Ast_3
Neu_1
Neu_2
Neu_3
Neu_4
Cell type
Zfp36
Fos
Id4
Id2
Mfap4
Neurod1
Hes6
Rnd2
Dcx
Mki67
Tubb6
Id3
Hes1
z−score
−2.5 0 2.5
C
Up-regulated
telencephalon development
regulation of cell projection organization
cell projection morphogenesis
neural precursor cell proliferation
central nervous system neuron differentiation
cerebral cortex cell migration
regulation of neuron differentiation
telencephalon regionalization
Down-regulated
cellular response to organic cyclic compound
response to wounding
extracellular matrix organization
positive regulation of cell motility
response to growth factor
mitotic cell cycle
response to nutrient
negative regulation of cell differentiation
-Log(P_value)
D
Hes1
Col11a1
Apoe
Aqp4
Gfap
Hes6
Neurod1
Prom1
Mfap4
Tubb3
max
min
E
Normalized accessibility (range 0 – 80)
Ast_3
Ast_2
Ast_1
ImA
ImA_ND1_hi
Neu_1
Neu_2
Neu_3
Neu_4
Gene
Peak
Aqp4
Gfap
FAM187A
Neurod2
Dcx
6505000 6515000 6525000
87850000 87860000
83320000
107425000 107475000
F
Accessibility
-3 kb
peak center
3 kb
G
NEUROD1_ChIP
Normalized accessibility
1kb
5kb
Gene
Cebpd
Fos
Anxa1
H
Neu_4
Neu_3
Neu_2
Neu_1
ImA_ND1_hi
ImA
Ast_1
Ast_2
Ast_3
Atf6, Elf2/4/5, Elk1/3/4, Etv1/2/3/4/5/6, Otx2, Xbp1, Irf2/3/4/5/7/8, Klf3/9/10/17, Pitx1/2/3, Rora/b/c, Bcl6, Erf/g, Rxra/b/g, Spi1/b/c,Sp2, Runx2/3, Nr2f1
Jun, Fos, Atf2/3/4/7, Cebpa/b/d/e/g/l, Batf, Bach1/2, Irf1/9, Klf2/4/5/6/11/14/15/16, Nfe2, Maf, Tead1/2/3/4, Sp1/3/4/8/9, Fosl1/2, Nfatc2/3/4, Rfx1/2/3/4/5, Stat1/3, Elf1/3,Hsf1/2, Junb/d
Neurod1/2, Neurog1/2, Olig1/2/3, Snai1/2/3, Id4, Ascl1, Atoh1/7, Tal1, Tcf3/4/12/21, Twist1
Dlx5/6, Emx1/2, Eomes, Msx1/2, Hes6, Lbx1/2, Gsx1/2, Smad2, Hoxd3/4/8/9, Hoxa1/2/4/5/6/7/10, Ctcf, Gata1/4/6, Pdx1, Hoxb2/3/4/5/6/7/8/13, Hoxc4/8, Lmx1a/b, Lhx1/2/5/6/9, Vsx1/2, Meis1/2/3, Evx1/2, Nkx6-1/2/3, Tbr1, Tlx2, Pax3/4/7, Prrx1/2, Tbx1/2/3/4/5/6/15/18/20/21, Pou2f1/2/3...
Motif score
-3 0 3
Pathway enrichment
DNA-templated transcription
cellular response to lipid
integrated stress response signaling
Th17 cell differentiation
regulation of neuron differentiation
CNS neuron differentiation
pattern specification process
sensory organ development
forebrain development

Figure S5

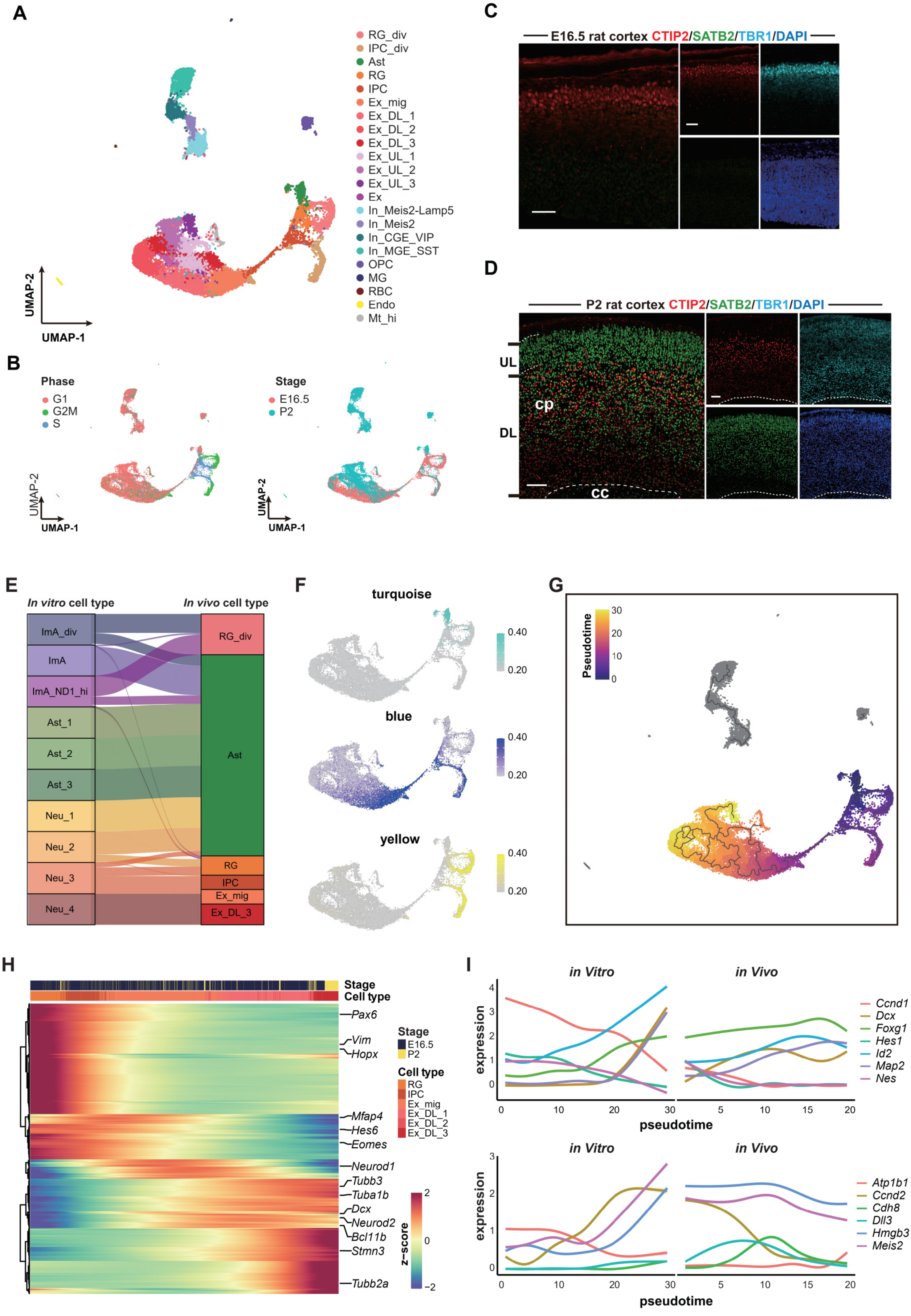
A
RG_div
IPC_div
Ast
RG
IPC
Ex_mig
Ex_DL_1
Ex_DL_2
Ex_DL_3
Ex_UL_1
Ex_UL_2
Ex_UL_3
Ex
In_Meis2-Lamp5
In_Meis2
In_CGE_VIP
In_MGE_SST
OPC
MG
RBC
Endo
Mt_hi
UMAP-1
UMAP-2
B
Phase
G1
G2M
S
Stage
E16.5
P2
C
E16.5 rat cortex CTIP2/SATB2/TBR1/DAPI
D
P2 rat cortex CTIP2/SATB2/TBR1/DAPI
UL
cp
DL
cc
E
In vitro cell type
In vivo cell type
ImA_div
ImA
ImA_ND1_hi
Ast_1
Ast_2
Ast_3
Neu_1
Neu_2
Neu_3
Neu_4
RG_div
Ast
RG
IPC
Ex_mig
Ex_DL_3
F
turquoise
blue
yellow
0.40
0.20
G
Pseudotime
H
Stage
Cell type
Pax6
Vim
Hopx
Mfap4
Hes6
Eomes
Neurod1
Tubb3
Tuba1b
Dcx
Neurod2
Bcl11b
Stmn3
Tubb2a
z-score
I
in Vitro
in Vivo
expression
pseudotime
Ccnd1
Dcx
Foxg1
Hes1
Id2
Map2
Nes
Atp1b1
Ccnd2
Cdh8
Dll3
Hmgb3
Meis2

Figure S6

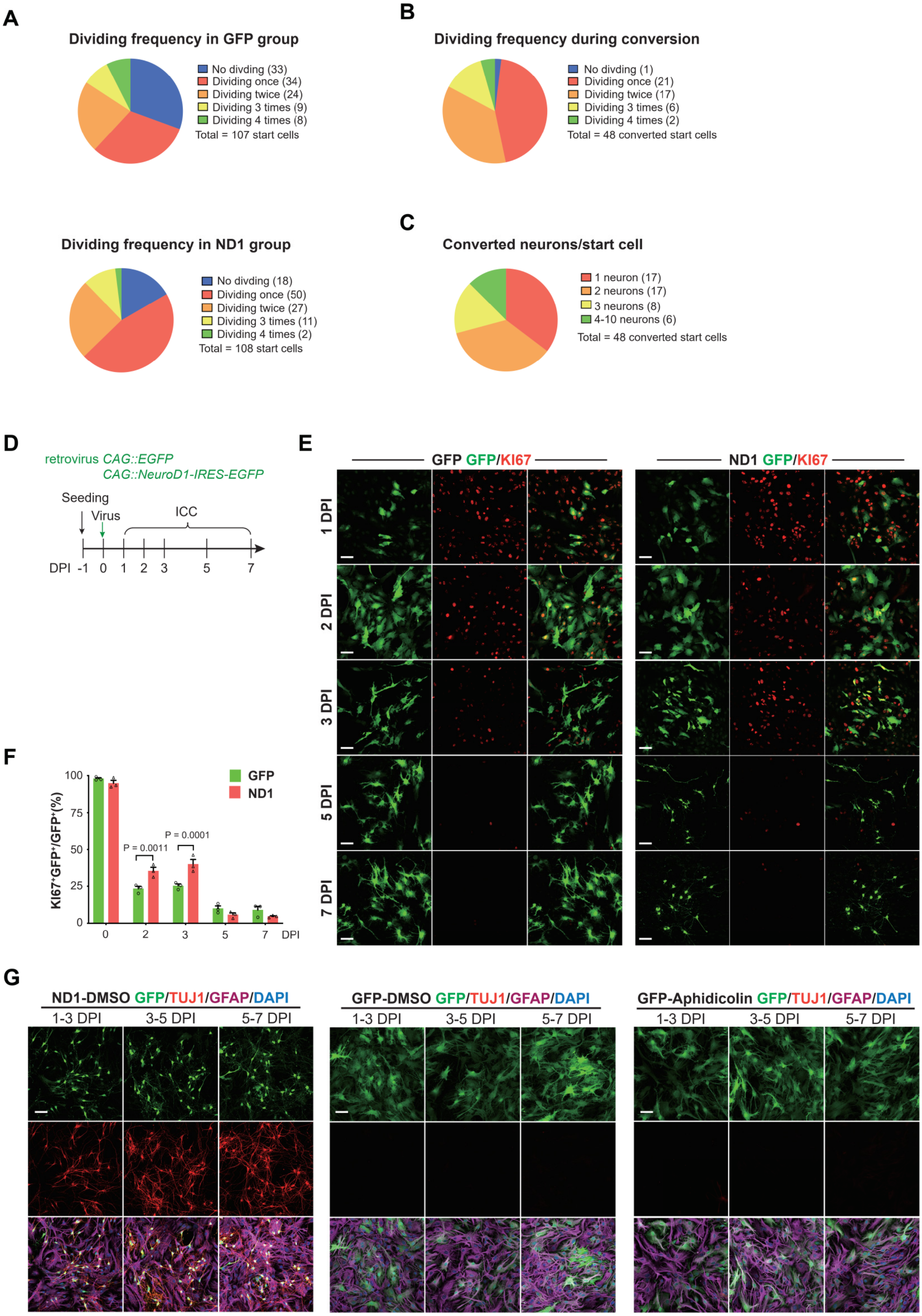

A
Dividing frequency in GFP group
No divding (33)
Dividing once (34)
Dividing twice (24)
Dividing 3 times (9)
Dividing 4 times (8)
Total = 107 start cells
Dividing frequency in ND1 group
No divding (18)
Dividing once (50)
Dividing twice (27)
Dividing 3 times (11)
Dividing 4 times (2)
Total = 108 start cells
B
Dividing frequency during conversion
No divding (1)
Dividing once (21)
Dividing twice (17)
Dividing 3 times (6)
Dividing 4 times (2)
Total = 48 converted start cells
C
Converted neurons/start cell
1 neuron (17)
2 neurons (17)
3 neurons (8)
4-10 neurons (6)
Total = 48 converted start cells
D
retrovirus CAG::EGFP
CAG::NeuroD1-IRES-EGFP
Seeding
Virus
ICC
DPI -1 0 1 2 3 5 7
E
GFP GFP/KI67
ND1 GFP/KI67
1 DPI
2 DPI
3 DPI
5 DPI
7 DPI
F
KI67+GFP+/GFP+(%)
GFP
ND1
P = 0.0011
P = 0.0001
100
75
50
25
0
0 2 3 5 7 DPI
G
ND1-DMSO GFP/TUJ1/GFAP/DAPI
GFP-DMSO GFP/TUJ1/GFAP/DAPI
GFP-Aphidicolin GFP/TUJ1/GFAP/DAPI
1-3 DPI
3-5 DPI
5-7 DPI

Figure S7

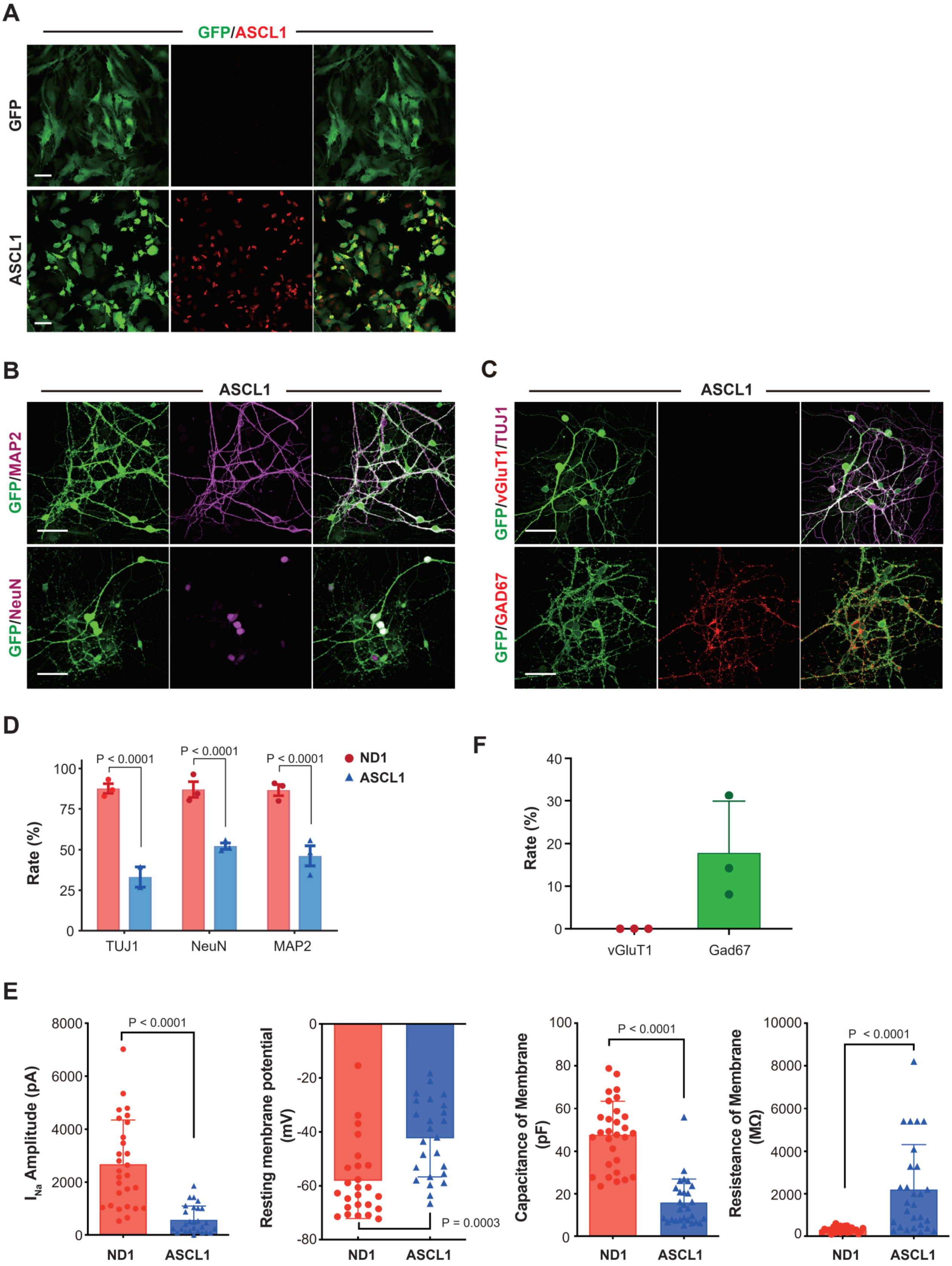

A
GFP/ASCL1
GFP
ASCL1
B
ASCL1
GFP/MAP2
GFP/NeuN
C
ASCL1
GFP/vGluT1/TUJ1
GFP/GAD67
D
Rate (%)
P < 0.0001
P < 0.0001
P < 0.0001
ND1
ASCL1
TUJ1
NeuN
MAP2
F
Rate (%)
vGluT1
Gad67
E
I Na Amplitude (pA)
P < 0.0001
ND1
ASCL1
Resting menbrane potential (mV)
P = 0.0003
ND1
ASCL1
Capacitance of Membrane (pF)
P < 0.0001
ND1
ASCL1
Resisteance of Membrane (MΩ)
P < 0.0001
ND1
ASCL1

Figure S8

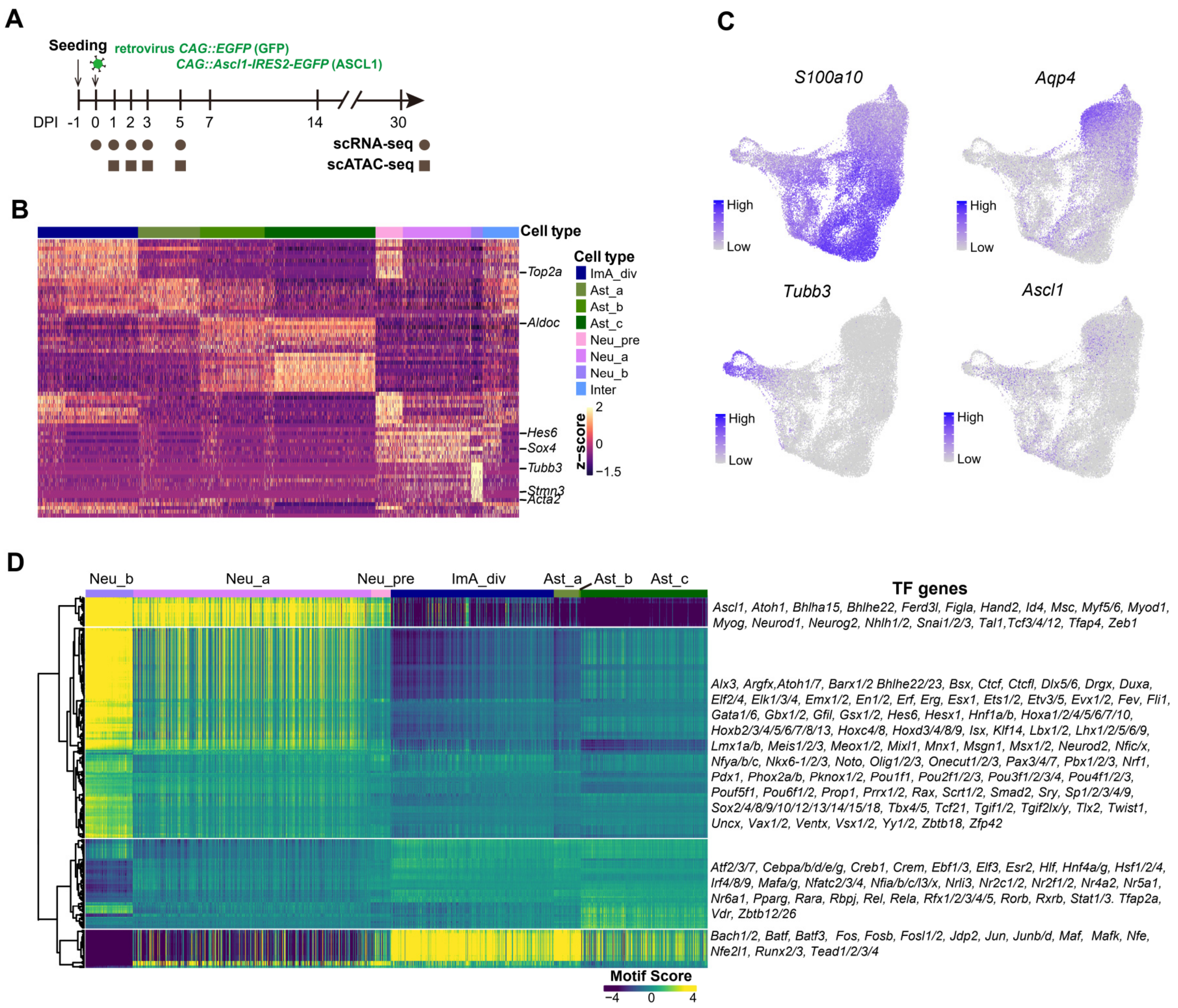

A
Seeding
retrovirus CAG::EGFP (GFP)
CAG::Ascl1-IRES2-EGFP (ASCL1)
DPI -1 0 1 2 3 5 7 14 30
scRNA-seq
scATAC-seq
B
Cell type
Top2a
Aldoc
Hes6
Sox4
Tubb3
Stmn3
Acta2
Cell type
ImA_div
Ast_a
Ast_b
Ast_c
Neu_pre
Neu_a
Neu_b
Inter
z-score
2
0
−1.5
C
S100a10
Aqp4
Tubb3
Ascl1
High
Low
D
Neu_b
Neu_a
Neu_pre
ImA_div
Ast_a
Ast_b
Ast_c
TF genes
Ascl1, Atoh1, Bhlha15, Bhlhe22, Ferd3l, Figla, Hand2, Id4, Msc, Myf5/6, Myod1, Myog, Neurod1, Neurog2, Nhlh1/2, Snai1/2/3, Tal1,Tcf3/4/12, Tfap4, Zeb1
Alx3, Argfx,Atoh1/7, Barx1/2 Bhlhe22/23, Bsx, Ctcf, Ctcfl, Dlx5/6, Drgx, Duxa, Elf2/4, Elk1/3/4, Emx1/2, En1/2, Erf, Erg, Esx1, Ets1/2, Etv3/5, Evx1/2, Fev, Fli1, Gata1/6, Gbx1/2, Gfil, Gsx1/2, Hes6, Hesx1, Hnf1a/b, Hoxa1/2/4/5/6/7/10, Hoxb2/3/4/5/6/7/8/13, Hoxc4/8, Hoxd3/4/8/9, Isx, Klf14, Lbx1/2, Lhx1/2/5/6/9, Lmx1a/b, Meis1/2/3, Meox1/2, Mixl1, Mnx1, Msgn1, Msx1/2, Neurod2, Nfic/x, Nfya/b/c, Nkx6-1/2/3, Noto, Olig1/2/3, Onecut1/2/3, Pax3/4/7, Pbx1/2/3, Nrf1, Pdx1, Phox2a/b, Pknox1/2, Pou1f1, Pou2f1/2/3, Pou3f1/2/3/4, Pou4f1/2/3, Pou5f1, Pou6f1/2, Prop1, Prrx1/2, Rax, Scrt1/2, Smad2, Sry, Sp1/2/3/4/9, Sox2/4/8/9/10/12/13/14/15/18, Tbx4/5, Tcf21, Tgif1/2, Tgif2lx/y, Tlx2, Twist1, Uncx, Vax1/2, Ventx, Vsx1/2, Yy1/2, Zbtb18, Zfp42
Atf2/3/7, Cebpa/b/d/e/g, Creb1, Crem, Ebf1/3, Elf3, Esr2, Hlf, Hnf4a/g, Hsf1/2/4, Irf4/8/9, Mafa/g, Nfatc2/3/4, Nfia/b/c/l3/x, Nrli3, Nr2c1/2, Nr2f1/2, Nr4a2, Nr5a1, Nr6a1, Pparg, Rara, Rbpj, Rel, Rela, Rfx1/2/3/4/5, Rorb, Rxrb, Stat1/3. Tfap2a, Vdr, Zbtb12/26
Bach1/2, Batf, Batf3, Fos, Fosb, Fosl1/2, Jdp2, Jun, Junb/d, Maf, Mafk, Nfe, Nfe2l1, Runx2/3, Tead1/2/3/4
Motif Score
−4
0
4

Figure S9

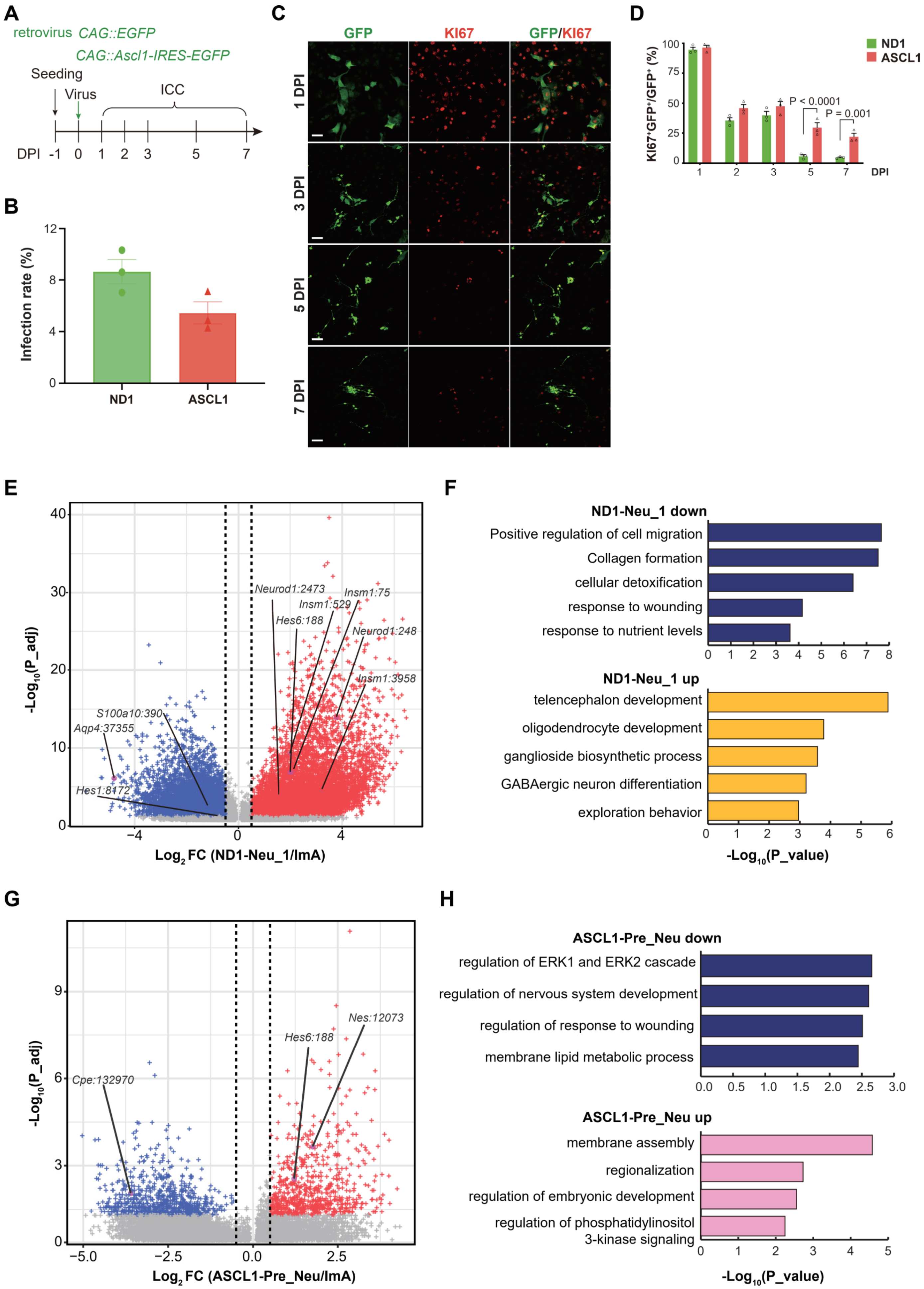
A
retrovirus CAG::EGFP
CAG::Ascl1-IRES-EGFP
Seeding
Virus
ICC
DPI -1 0 1 2 3 5 7
B
Infection rate (%)
ND1
ASCL1
C
GFP
KI67
GFP/KI67
1 DPI
3 DPI
5 DPI
7 DPI
D
KI67+GFP+/GFP+ (%)
ND1
ASCL1
P < 0.0001
P = 0.001
DPI
E
Neurod1:2473
Insm1:75
Insm1:529
Hes6:188
Neurod1:248
Insm1:3958
S100a10:390
Aqp4:37355
Hes1:8172
-Log10(P_adj)
Log2 FC (ND1-Neu_1/ImA)
F
ND1-Neu_1 down
Positive regulation of cell migration
Collagen formation
cellular detoxification
response to wounding
response to nutrient levels
ND1-Neu_1 up
telencephalon development
oligodendrocyte development
ganglioside biosynthetic process
GABAergic neuron differentiation
exploration behavior
-Log10(P_value)
G
Nes:12073
Hes6:188
Cpe:132970
-Log10(P_adj)
Log2 FC (ASCL1-Pre_Neu/ImA)
H
ASCL1-Pre_Neu down
regulation of ERK1 and ERK2 cascade
regulation of nervous system development
regulation of response to wounding
membrane lipid metabolic process
ASCL1-Pre_Neu up
membrane assembly
regionalization
regulation of embryonic development
regulation of phosphatidylinositol
3-kinase signaling
-Log10(P_value)

### Supplementary Figure Legends

**Figure S1 (related to Figure 1). ND1-induced *in vitro* astrocyte-to-neuron conversion**

(A-B) Representative images (A) and quantitative analysis (B) show that the majority of primary cells isolated from neonatal rat cortices at passage 5 (P5) were astrocytes. GFAP (green), S100B (green), VIMENTIN (green) are markers for astrocytes. TUJ1 (green) is maker for neurons. GFAP- and OLIG2+ is marker for oligodendrocytes. Scale bars = 100 μm. (C-D) Representative images (C) and quantitative analysis (D) show that the majority of infected cells are astrocytes. Scale bars = 50 μm.

(E) Representative images reveal the expression NEUROD1 (red) in the infected cells (GFP) at 3 DPI. DPI: days post infection. Scale bar = 50 μm.

(F) Representative images show the gradual morphology change and expression of neuronal marker TUJ1(red) in the ND1 group. Scale bar = 50 μm.

(G) Representative images show the expression of markers for immature (DCX in purple) and mature (MAP2 in red) neurons at 7 and 14 DPI. Scale bar = 50 μm.

(H-K) Representative images (H-I) and quantitation (J-K) reveal the high neuronal conversion rate and that most converted neurons are cortical deeper-layer exciting neurons. Scale bars = 50 μm (H) and 20 μm (I).

**Figure S2 (related to Figure 1). ND1-converted neurons display similar electrophysiological features to those of primary neurons**

(A-B) Representative traces (A) and quantitative analysis (B) show comparable sodium currents of ND1-converted neurons to that of primary neurons isolated from E16.5 rat cortices.

(C-E) Quantitation reveals similar resting membrane potential (C), capacitance of membrane (D) and resistance of membrane (E) between ND1-converted neurons and primary neurons.

(F-H) Representative traces (F) and quantitative analysis (G-H) show the modes of action potential of the ND1-converted neurons, as well as the similarity to the primary neurons.

(I-K) Representative traces (I) and quantitative analysis (J-K) reveal that the ND1-converted neurons form mainly excitable neural circuits that share comparable parameters to those of primary neurons.

**Figure S3 (related to Figure 2). Regulating network of ND1-induced astrocyte to neuron reprogramming**

(A) Pseudotime results by Monocle2. The expression levels of astrocyte gene Aqp4 and neuronal gene Tubb3 are shown on the right to confirm the branch identity.

(B) Relative expression of identical genes along the two branches.

(C) WGCNA modules clustered based on all cells in ND1 and GFP group.

(D) The expression pattern of each module on the UMAP.

(E) The gene network within each module.

(F) GO results of genes in each module.

**Figure S4 (related to Figure 3). Multiomics analysis of the ND1-induced astrocyte to neuron reprogramming**

(A) Annotation of peaks obtained by NEUROD1 ChIP-seq.

(B) Expression patterns of the NEUROD1 target genes.

(C) GO results of genes directly upregulated and downregulated by ND1.

(D) Gene score of cell type markers.

(E) The genome browser view showing the chromatin accessibility signal.

(F) The accessibility of NEUROD1 ChIP peaks in each cell type.

(G) The genome browser view showing the NEUROD1 ChIP signal and chromatin accessibility signal.

(H) Heatmap showing the activity values of the TFs predicted important in

ND1-induced immature astrocytes to neuronal reprogramming. The TFs are clustered in to four groups and the GO terms of each group are shown in the right.

**Figure S5 (related to Figure 4). ScRNA-seq analyses of *in vivo* neurogenesis**

(A) UMAP showing the cell types captured in E16.5 and P2 rat cortice.

(B) The cell cycle states and group information of each cell.

(C-D) Representative images show the composition of rat cortical neurons at E16.5 (C) and P2 (D). Scale bars = 50 μm (C) and 100 μm (D). CTIP2 (red) and TRB1 (turquoise) are markers representing deep-layer neurons. SATB2 (green) is marker representing upper-layer neurons. DAPI (blue) represents nuclei. UL: upper layer; DL: deep layer; cp: cortical plate; cc: corpus callosum.

(E) The congruent relationship of *in vitro* and *in vivo* cell types.

(F) The expression pattern of three WGCNA modules representing astrocyte, neuron and progenitor respectively on the *in vivo* cells.

(G) Pseudotime score of the *in vivo* cells on UMAP.

(H) Heatmap showing the expression of *in vivo* neurogenesis genes along pseudotime.

(I, J) Expression patterns of representative consistent genes (I) and inconsistent genes (J) along *in vitro* and *in vivo* pseudotime.

**Figure S6 (related to Figure 5). Cell dividing is required for ND1-induced neuronal reprogramming**

(A) Quantitation of dividing frequency of the start cells in the GFP (upper) and ND1 (lower) group based on the time-lapse live image data.

(**B**) Quantitation of dividing frequency of the start cells successfully converted to neurons.

(C) Quantitation of the number of converted neurons per start cell in ND1 group.

(D) Schematic of the experiment design to investigate the proliferating cells during ND1-induced neuronal reprogramming.

(E-F) Representative images (E) and quantitation (F) showing more $KI67^+$ (red) cells in the infected cells ($GFP^+$, green) in the early stage of neuronal reprogramming (2,3 DPI). Scale bars = 50 μm.

(G) Representative images showing that DMSO (vehicle of aphidicolin) has little effect on ND1-induced neuronal reprogramming. GFP (green) represents infected cells. TUJ1 (red) represents neurons. GFAP (purple) represents astrocytes. DAPI (blue) represents nuclei. Scale bars = 50 μm.

**Figure S7 (related to Figure 6). ASCL1-induced *in vitro* neuronal reprogramming**

(A) Representative images showing the expression of ASCL1 (red) at 3 DPI. Scale bar = 50 μm.

(B-F) Representative images (B-C) and quantitation (D-F) show that ASCL1-induced neuronal reprogramming is less efficient than that induced by ND1, and the converted neurons are inhibitory neurons. GAD67 (red) represents inhibitory neurons. Scale bars = 50 μm.

(E) Quantitative analysis revealing ASCL1-induced neurons are immature in electrophysiological features compared with ND1-converted neurons.

**Figure S8 (related to Figure 6). Single-cell analyses of ASCL1-induced AtN conversion**

(A) Schematic of the *in vitro* ASCL1-induced neuronal reprogramming system and sequencing strategies.

(B) Heatmap showing the expression patterns of cell type DEGs.

(C) The expression levels of indicated markers.

(D) Heatmap showing the activity values of the TFs predicted important in ASCL1-induced immature astrocytes to neuronal reprogramming.

**Figure S9 (related to Figure 7). Different molecular changes induced by ND1 and ASCL1**

(A) Schematic of the experiment design to investigate the proliferating cells during ASCL1-induced neuronal reprogramming.

(B-E) Representative images (C) and quantitation (B,D) showing similar infection rate between ND1 and ASCL1 group, and that more $KI67^{+}$ (red) infected cells ($GFP^{+}$, green) in the ASCL1 group in relatively late stage of neuronal reprogramming (5,7 DPI). Scale bars = 50 μm.

(E) The DEGs between ND1_Neu1 and ImA.

(F) Representative GO terms of DEGs between ND1_Neu1 and ImA.

(G) The DEGs between ASCL1_Pre and ImA.

(H) Representative GO terms of DEGs between ASCL1_Pre and ImA.
